# Functional brain segmentation using inter-subject correlation in fMRI

**DOI:** 10.1101/057620

**Authors:** Jukka-Pekka Kauppi, Juha Pajula, Jari Niemi, Riitta Hari, Jussi Tohka

## Abstract

The human brain continuously processes massive amounts of rich sensory information. To better understand such highly complex brain processes, modern neuroimaging studies are increasingly utilizing experimental setups that better mimic daily-life situations. We propose a new exploratory data-analysis approach, functional segmentation intersubject correlation analysis (FuSeISC), to facilitate the analysis of functional magnetic resonance (fMRI) data sets collected in these experiments. The method provides a new type of functional segmentation of brain areas, not only characterizing areas that display similar processing across subjects but also areas in which processing across subjects is highly variable.

We tested FuSeISC using fMRI data sets collected during traditional block-design stimuli (37 subjects) as well as naturalistic auditory narratives (19 subjects). The method identified spatially local and/or bilaterally symmetric clusters in several cortical areas, many of which are known to be processing the types of stimuli used in the experiments. The method is not only prominent for spatial exploration of large fMRI data sets obtained using naturalistic stimuli, but has other potential applications such as generation of a functional brain atlases including both lower-and higher-order processing areas.

Finally, as a part of FuSeISC, we propose a criterion-based sparsification of the shared nearest-neighbor graph for detecting clusters in noisy data. In our tests with synthetic data, this technique was superior to well-known clustering methods, such as Ward's method, affinity propagation and K-means++.

## 1. Introduction

Traditionally, neuroimaging studies have utilized highly controlled and simplified experimental setups to study human brain function. While these studies have been, and continue to be, extremely informative, the applied simplified stimuli do not resemble situations of daily life, where the brain continuously receives massive amounts of rich sensory information. In recent years, attempts have been made to conduct more naturalistic experiments that better mimic daily life and thus should help to understand complex brain processes.

While the amount of complex neuroimaging data sets collected in naturalistic experiments is increasing, a major bottleneck remains to be the lack of proper analysis methods. So far, one of the most promising approaches to analyze such complex functional magnetic resonance imaging (fMRI) data sets is *inter-subject correlation (ISC) analysis* (Hasson et al., 2004), applied to fMRI data sets collected using naturalistic stimuli, such as movies/video (Hasson et al., 2004, Golland et al., 2007, Nummenmaa et al., 2012a, Reason et al., 2016) and music (Trost et al., 2015, Abrams et al., 2013). ISC-based analysis is conceptually simple, involving voxel-wise computations of correlation coefficients between time series of all subjects. Once the correlation coefficients have been computed across all participants exposed to the identical time-varying stimulus sequence, the subject-pair-wise correlation coefficients for each voxel can be averaged and subsequently thresholded to obtain brain maps indicating which regions exhibit considerable ISC during the stimulation (Wilson et al., 2008, Kauppi et al., 2010b). A major strength of the ISC-based analysis is that it can detect activated brain areas without modeling the expected hemodynamic responses (Pajula et al., 2012).

Despite its benefits, the existing ISC-based analysis has limitations. For example, it typically provides voxel-wise information about the extent of the ISCs during the whole fMRI time series of interest. For longer time series, ISC can be computed in several shorter time windows (see e.g. Nummenmaa et al. (2012b)), but there exists no standard procedure how to integrate ISC information across the time windows.

In any case, integrating ISC information across voxels and time frames of interest may provide new insights into functional architecture of the human brain. More specifically, each voxel can be characterized by a pattern of ISC features, describing how extensively a voxel is co-activated during different stimuli of interest. For instance, out of five different video/audio clips, a voxel may not show any ISC during two clips but may exhibit very high ISC during one clip and moderate ISC during the remaining two clips. It is plausible to assume that some voxels share a highly similar pattern of ISC features whereas some other voxels do not, meaning that voxels can be organized into distinct clusters on the basis of these features. Thus, to better understand the functional organization of the human brain during processing of complex stimuli, we propose formation of subject-pair-wise averaged ISC features from specific time series of interest, and clustering them across the brain.

Another limitation of the conventional ISC-based mapping is that it assumes similar brain mechanisms across subjects.^1^ It is, however, well known that individuals process identical sensory information more or less differently, especially in higher-order brain areas that are strongly involved in situations of daily life (Hasson et al., 2010). Therefore, a conventional ISC approach based on the averaging of correlation coefficients across all pairs of subjects may find high ISC values in sensory projection areas but may completely lose ISC in higher-order brain areas due to high inter-subject variability (Kauppi et al., 2010a). Consequently, averaging across subjects abolishes signs of active processing in such important brain areas.

To better understand the functions of different brain areas, we incorporated into our analysis subject-pair-wise ISC variability in addition to traditional averaging of ISC features. It is likely that brain areas of high average ISC together with relatively low ISC variability mostly reflect sensory processing that is expected to be most coherent across subjects. On the other hand, areas with relatively high ISC variability may reveal activations with higher inter-individual differences. Note that, in contrast to our interpretation, inter-subject variation of signal strengths is in neuroimaging data traditionally considered as noise. However, recent studies show that individual variability provides meaningful information that can elucidate complex brain processes and brain development (Mueller et al., 2012, Zilles and Amunts, 2013, Boldt et al., 2014, Gopal et al., 2016).

We call our entire method, which combines ISC-based feature extraction and clustering, functional segmentation ISC analysis (FuSeISC). The features are extracted from multiple subjects and multiple fMRI time series of interest. The fMRI time series can be selected either from separate experiments, separate runs within the same experiment, or from selected time intervals of a longer fMRI experiment (for example, corresponding to the scenes of a movie). Due to both local and distributed brain processes, it is likely that some of the clusters found in the “ISC feature space” are spatially local whereas others are more widely spread. Therefore, we do not apply to the segmentation any anatomical constraints. The FuSeISC method described in this paper won the Study Forrest Real Life Cognition Challenge^2^ (Hanke et al., 2014) where the goal was to introduce novel analysis methods for complex fMRI data sets acquired under naturalistic stimulation. Here, we present the details of the algorithm and validate the technique more thoroughly with different data sets. FuSeISC has been integrated to the ISC toolbox (Kauppi et al., 2014) and is freely available at https://www.nitrc.org/projects/isc-toolbox/.

We have previously presented clustering of ISC matrices (Kauppi et al., 2010a) to analyze how subject-pair-wise ISCs are distributed across brain areas during a complex stimulus time course. FuSeISC notably extends this approach by capturing spatiotemporal variation in ISCs as it utilizes a number of shorter time series instead of a single time course. Another major difference is that in FuSeISC, we cluster features describing the summary statistics (mean, variability) of the ISC matrices instead of the entire matrices. This procedure is important because the number of subject-pair-wise ISCs (dimensionality) increases rapidly together with the number of time series and subjects. In FuSeISC, we also replace a random initialization approach used in Kauppi et al. (2010a) with a new algorithm which provides more reliable initial estimates of cluster centroids. Finally, we replace a *K*-means algorithm used in Kauppi et al. (2010a) with model-based clustering which allows finding clusters with more complex covariance structures.

## 2. Materials

### 2.1. ICBM Functional Reference Battery data

The fMRI data collected during Functional Reference Battery (FRB) tasks developed by the International Consortium for Human Brain Mapping (ICBM) (Mazziotta et al., 2001) were used for the evaluation of the method and for the construction of the simulated data set described in the next subsection. The block-design FRB tasks are a set of behavioral tasks designed to produce reliable across-subjects functional landmarks in brain-imaging data, and the data sets as such are ideal for the validation of functional segmentation methods. We have previously used the same data for other experiments. For details of the data and experiments, see Pajula et al. (2012) and Pajula and Tohka (2014), but, for convenience, we provide a short description here.

The FRB fMRI data were obtained from the ICBM database in the Image Data Archive of the Laboratory of Neuro Imaging. The ICBM project (Principal Investigator John Mazziotta, M.D., University of California, Los Angeles) is supported by the National Institute of Biomedical Imaging and BioEngineering. ICBM is the result of efforts of co-investigators from UCLA, Montreal Neurological Institute, University of Texas at San Antonio, and the Institute of Medicine, Jülich/Heinrich Heine University, Düsseldorf, Germany.

The dataset used earlier by Pajula et al. (2012) included fMRIs of 41 right-handed subjects whose fMRI had been measured during five FRB tasks: (1) auditory naming (AN) task where the subject silently named objects that were verbally described; (2) external ordering (EO) task where the subject, after a delay period (and thus relying on working memory), kept track of the abstract designs on the screen; (3) hand imitation (HA) task where the subject was instructed to imitate the presented hand configuration with his right hand; (4) oculomotor (OM) task where the subject made saccades to target locations; and (5) verb generation (VG) task where the subject generated a verb that corresponded to an object presented on the screen. For detailed definitions of the five FRB tasks, see the FRB software package^3^ and Pajula et al. (2012). Pajula et al. (2012) discarded four subjects during the pre-screening phase because of poor data quality in at least one task in the battery. Thus, the final data set consisted of measurements from 37 healthy right-handed subjects (19 men and 18 women; mean age 28.2 years, range 20–36).

In addition to the original ICBM data set, we also investigated the reproducibility of the FuSeISC results with two ICBM data sets consisting of different subjects. For this purpose, we selected altogether 74 subjects from the ICBM database by widening the original age range of the subjects (the ages of the subjects in this new data set were between 21 and 55 years). The data set was then split into two comparable sets both consisting of 37 subjects. Furthermore, we investigated the effect of the number of subjects on the results by forming four additional data sets from the whole 74 subject set: An age-matched pair of data sets with 25 subjects and another age-matched pair of data sets with 15 subjects. Table 1 lists the details of the data sets.

**Table 1:** Description of ICBM data sets used to compare FuSeISC clustering with different sets of subjects. Data sets were balanced to have close to equal number of male and female subjects as well as similar age range and mean age. A single subject appeared only in one of the two data sets (#1 or #2). First row (ICBM37ORIG) is the data set from Pajula et al. (2012).

| Data set | Age range | Mean age | # Male | # Female |
| --- | --- | --- | --- | --- |
| ICBM37ORIG | 20–36 | 28.2 | 19 | 18 |
| ICBM37#1 | 21–55 | 37.6 | 19 | 18 |
| ICBM37#2 | 21–54 | 37.4 | 20 | 17 |
| ICBM25#1 | 21–54 | 37.2 | 13 | 12 |
| ICBM25#2 | 21–55 | 38.3 | 13 | 12 |
| ICBM15#1 | 21–53 | 35.9 | 8 | 7 |
| ICBM15#2 | 21–53 | 36.9 | 8 | 7 |

The functional fMRI data were collected with a 3 T Siemens Allegra FMRI scanner and the anatomical T1 weighted MRI data with a 1.5 T Siemens Sonata scanner. The TR/TE times for the functional data were 4 s/32 ms, flip angle 90°, pixel spacing 2 mm and slice thickness 2 mm. There were 12 blocks of 7 volumes per task (6 ‘off-on’ blocks) and 3 volumes at the beginning of the run to wait for magnetisation stabilisation, which were removed during the preprocessing. The total lengths of the time series in the analysis were 84 volumes (with the total duration of 5 min 36 s). The acquisition parameters for the anatomical T1 data were 1.1 s/4.38 ms, 15°, 1 mm and 1 mm, correspondingly. Preprocessing was performed as described in Pajula et al. (2012) by a standard FSL preprocessing pipeline including Gaussian 5-mm full width at half maximum (FWHM) spatial filtering.

### 2.2. Simulated data

We generated synthetic fMRI data sets based on the ICBM data described above. Similarly to the experimental ICBM data, the simulated data consisted of five FRB tasks (AN, EO, HA, OM, and VG) from 37 subjects. The purpose of simulated data was to validate the functional segmentation method quantitatively when the true functional segmentation is fully known.

In the simulated data sets and for each task separately, every voxel was defined either as “activated” or “non-activated”. Thus, any voxel was characterized by a 5-element binary vector creating 2^5^ = 32 distinct functional segments. Voxels were selected as “activated” according to the binarized statistical maps of the GLM analysis performed for the empirical ICBM data sets in Pajula et al. (2012) (thresholded at voxel-wise false discovery rate (FDR) corrected threshold *q* = 0.001). A simulated hemodynamic signal was included in the time series of the activated voxels; the signal was identical to the one used as a model in the GLM analysis of the data (see Pajula et al. (2012)), i.e., a boxcar convolved with a canonical hemodynamic response function (HRF). These signals were then corrupted by pink 1/f noise which was generated according to Smith (2012). Signal-to-noise-ratio (SNR) was 0.02, which was quantified on the basis of the boxcar function *before* the convolution with the canonical HRF. All brain areas outside the activated regions contained only noise.

The generation procedure was identical for every 37 simulated data sets and FRB tasks. We ignored anatomical and effect size variations between the subjects. Moreover, since the original empirical data sets were registered to the MNI-152 coordinate space, we did not perform registration or motion correction as preprocessing. The preprocessing only included Gaussian 5-mm FWHM spatial filtering.

### 2.3. StudyForrest data

To demonstrate the performance of the FuSeISC method with naturalistic stimulation, we analyzed fMRI data sets of 19 subjects provided by the organization committee of the StudyForrest project and data challenge. The details of the experiment, data collection and preprocessing are provided by Hanke et al. (2014). In brief, the participants listened to a German sound track (Koop, Michalski, Beckmann, Meinhardt & Benecke, produced by Bayrischer Rundfunk, 2009), of the movie “Forrest Gump” (R. Zemeckis, Paramount Pictures, 1994) as broadcast as an additional audio track for visually-impaired listeners on Swiss public television.

The auditory content was largely identical to the dubbed German sound track of the movie, including the original dialogues and environmental sounds, but added by interspersed narrations by a male speaker who described the visual contents of the scenes. As detailed by Hanke et al. (2014), the participants listened to the movie sounds using custom-built in-ear headphones designed to maximize comfort during the scanning. T2-weighted echo-planar images (gradient-echo, 2-s TR, 22-ms echo time, 0.78-ms echo spacing, generalized autocalibrating partially parallel acquisition (GRAPPA)) were acquired during stimulation using a whole-body 7 T Siemens MAGNETOM scanner. Altogether 36 axial slices (thickness 1.4 mm, 1.4 mm × 1.4 mm in-plane resolution, 224-mm field-of-view, anterior-to-posterior phase encoding direction) with 10% interslice gap were recorded in ascending order. Slices were oriented to include the ventral portions of frontal and occipital cortex while minimizing the intersection with the eyeballs. Note that the brain coverage of the scans was limited due to the high scan resolution (Hanke et al., 2014).

The entire data set consisted of 8 runs (about 15 min each) for each subject from which we selected sound segments for our analysis. We selected five attractive sound segments, because we noted that they had created more buzz in Internet movie forums than the other scenes of the movie. We ranked the attractiveness of the clips based on an Internet survey of the corresponding video clips (that the subjects did not see) on online video services such as YouTube and movie-discussion forums. Table 2 lists the time points used to create the five clips. The exact data set for the analysis was extracted from the original preprocessed linear anatomical alignment set of the StudyForrest data. In addition to preprocessing performed by the providers of StudyForrest data (Hanke et al., 2014), we included Gaussian spatial filtering with the isotropic 3-mm FWHM kernel.

**Table 2:** Time points (in fMRI volumes) of the audio clips used in the analysis of StudyForrest data. Clip 2 has data from two acquisition sessions.

| Clip | Run | Start | Stop | Length | Description |
| --- | --- | --- | --- | --- | --- |
| Clip 0 | 1 | 1 | 50 | 50 | Feather flies and actors are described |
| Clip 1 | 2 | 1 | 50 | 50 | Scene with 'Run, Forrest, Run' cry |
| Clip 2 | 2 | 342 | 441 | 55 | Scene where Bubba and Forrest discuss about shrimps and how to cook them |
|  | 3 | 1 | 45 |  |  |
| Clip 3 | 6 | 158 | 404 | 246 | Forrest runs across the USA from coast to coast |
| Clip 4 | 7 | 46 | 107 | 51 | Next to the Jenny's bed, Forrest tells about his adventures, and at the end of the scene Jenny dies |

### 2.4. Resting-state fMRI data

In addition to stimulus-related fMRI data, we applied the FuSeISC method to resting-state fMRI (rfMRI) data of 38 randomly selected, unrelated subjects from the Human Connectome Project WU-Minn HCP Data -900+ 7 T data set (Essen et al., 2012). The data set included 17 males and 21 females with ages between 22 and 35 years. The data were pre-processed (Glasser et al., 2013) and co-registered by the Human Connectome Project (Marcus et al., 2011) non-linearly to a common MNI-152 space. For the data-acquisition protocol, see Essen et al. (2012). The first resting-state session (REST1) with the left-to-right scanning protocol (LR) was divided into 5 clips with 140 time points each. The total length of the session was 1200 time points. The first 10 time points as well as 10 time points between each clip were discarded. FuSeISC was then run for these five rfMRI clips.

## 3. Methods

The FuSeISC method consists of two main steps:

1. **Feature extraction** (Section 3.1): Given *M* fMRI time series of *N* subjects, 2*M* ISC-based features are extracted for each voxel, as illustrated in Fig. 1.
2. **Clustering** (Section 3.2): Feature vectors of the voxels are clustered to form the functional segmentation of the brain.

These steps, together with the performance-evaluation metrics, will be described next.

### 3.1. Feature extraction

Functional segmentation has been typically performed individually for each subject, based on the individual fMRI time series, and the individual clustering results have been combined in a subsequent stage to form group-level cluster maps (see, e.g., van den Heuvel et al. (2008)). We propose a different approach in which information is directly integrated across subjects by computing subject-pair-wise ISCs from multiple temporally distinct time series and extracting features from them. Two ISC features—the mean and the variability of pair-wise correlations—are extracted from the selected time series. They provide complementary information about processing in different brain regions.

Features were extracted separately for each voxel of the brain using the ISC toolbox (Kauppi et al., 2014), as described in Figure 1. For each of *M* time series, we computed correlation coefficients between the time series of all subject pairs, leading to *N × N* ISC matrix for each time series, where *N* is the number of subjects. For instance, the fMRI data sets of the Forrest study were divided into *M* = 5 distinct time series, corresponding to the five scenes of interest (see Section 2.3 on how the most interesting scenes were selected). The ISC features were computed based on the ISC matrices. First, the means of subject-pair-wise correlation coefficients, i.e., the *mean ISC features*, were computed for each time series *m* and for each voxel (a voxel index is omitted for clarity):

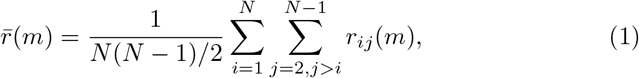
for *m* = 1, 2*,…, M*. Here, 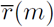 denotes a group-level ISC in a given voxel for time series *m* and *r*_*ij*_ (*m*) is the correlation coefficient between *m*th fMRI timecourses of subjects *i* and *j*. Note that because *r*_*ii*_(*m*) = 1 and *r*_*ij*_ (*m*) = *r*_*ji*_(*m*), it is sufficient to compute correlation coefficients across *N* (*N −* 1)*/*2 subject pairs (instead of *N*^2^ pairs) (Kauppi et al., 2014).

**Figure 1:**
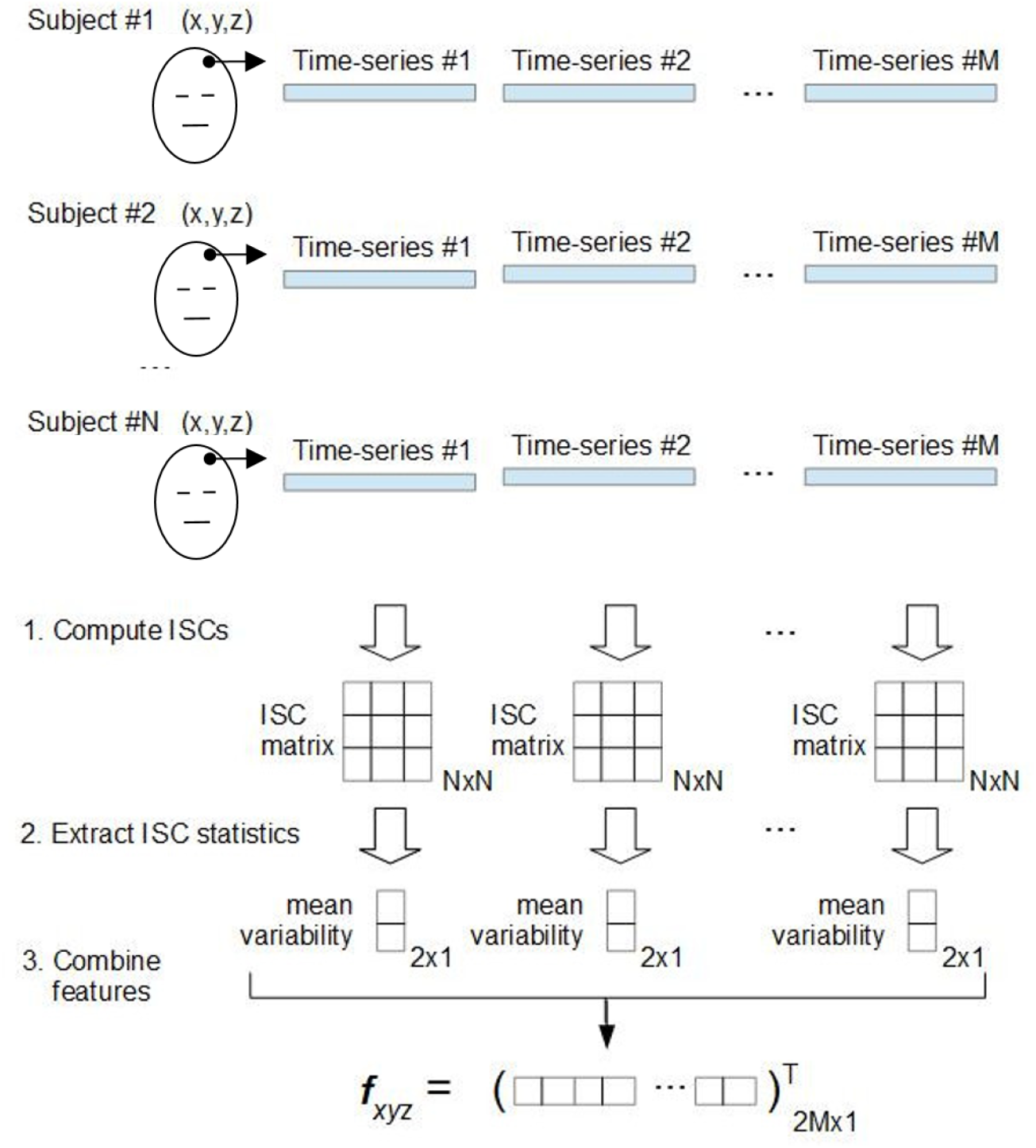
Illustration of the feature extraction in FuSeISC for one arbitrary voxel located at coordinate (x,y,z). At first, *M* ISC matrices are independently computed based on the fMRI time series of *N* subjects. In our study, the total number of time series was *M* = 5, corresponding to the total number of tasks (ICBM data) or movie clips (StudyForrest data) of interest. From each *N × N* ISC matrix, mean and variability are extracted using the Jackknife procedure. These two features are stacked into a single feature vector *f*_*xyz*_, whose dimension is 2*M*. This procedure is repeated for each brain voxel to obtain altogether 228,483 and 449,612 feature vectors for cluster analysis, corresponding to the ICBM and StudyForrest data, respectively.

We computed *ISC variability features* using a leave-one-subject-out Jackknife procedure, similar to that applied by Pajula and Tohka (2014). More specifically, we first computed the mean ISC values so that each subject was left out from the original sample one at a time. This procedure corresponds to the computation of the *N* mean ISC values, called *pseudovalues*, for *i* = 1, 2,…, *N*, so that *i*th row and *i*th column in the ISC matrix are left out one at a time. The Jackknife standard-error estimate was then computed as standard deviation of the pseudovalues multiplied by 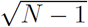. With simple algebraic manipulation, it can be shown that this procedure corresponds to computing

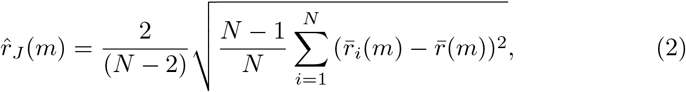
where 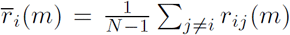. The Jackknife technique was preferred over the bootstrap due to a heavier computational burden associated with the bootstrap. Finally, the mean and variability features were combined into the feature vector

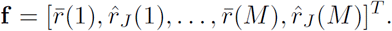

After feature extraction, we have one instance of the feature vector **f** for each voxel. The supporting idea in the above feature-extraction scheme is that voxels showing similar mean and variability statistics in ISCs for each time series of interest belong to the same functional segment. This way, the brain is divided into different functional regions on the basis of ISC features. Because the time series of interest have different characteristics in ISCs, it is likely that clustering reveals multiple brain areas, each constructed on the basis of a specific pattern of ISC mean and variability features. The number of time features (twice the number of time series *M*) should be much smaller than the number of voxels. No assumptions are made about the relationship between the number of subjects and the number of time series (i.e. *N > M* or *M ≥ N*). However, the more subjects we have, the less noisy are the features.

### 3.2. Robust algorithm for functional segmentation Gaussian mixture model

After the feature extraction, we learned a Gaussian mixture model (GMM) to cluster the ISC features. GMM provides a principled way of performing the functional segmentation under the assumption that the ISC features form clusters which follow a Gaussian distribution. Importantly, we did not impose any spatial constraints on our model, meaning that functional segments need not be spatially local but can consist of several spatially disjoint “subclusters”. The model is given by (McLachlan and Peel, 2000):

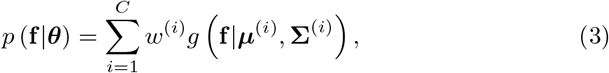
where *C* is the total number of clusters, 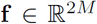 feature vector described in the previous section, ***θ*** denotes all the parameters of the model, *w*^(*i*)^ *∈* [0, 1], 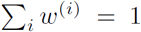 are mixture weight parameters, and 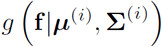 are multivariate Gaussian component densities with the mean ***µ***^(*i*)^ and the covariance 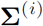. Because a multivariate Gaussian distribution can be fully described by its mean and covariance matrix, the unknown parameters of the GMM are 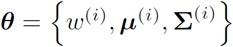, for *i* = 1, 2,…, *C*. The elements of 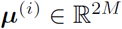 are given 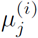 and the elements of 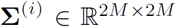 are given by 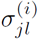. Note that the mean vector of each cluster ***µ***^(*i*)^ characterizes the cluster in terms of the original mean ISC and variability ISC features. We estimated the maximum likelihood solutions for these parameters using the expectation maximization (EM) algorithm (Xu and Jordan, 1996, McLachlan and Peel, 2000) implemented in the Statistics Toolbox of the Matlab.

#### Finding initial model

A major difficulty with the GMM-based clustering is that the quality of the clustering is highly dependent on a selected initial model (Fraley and Raftery, 2002, Figueiredo and Jain, 2002): if the mean vectors of the Gaussian components are not initially near the true cluster mean values, the EM algorithm converges towards a suboptimal solution and easily misses interesting clusters in the data.^4^ Another problem is that the total number of clusters *C* in the GMM is hard to determine because well-known model-selection criteria, such as the Bayesian information criterion (BIC), tend to overestimate the total number of clusters in complex fMRI data sets (Thirion et al., 2014).

To overcome these problems, we propose restricting a set of initial candidate models *a priori* to meaningful ones based on local structures in the data. Besides accuracy, prerequisites for the algorithm are computational and memory efficiency, because we run segmentation across all the brain voxels (the number of brain voxels was 228,483 for the ICBM data and 449,612 for the StudyForrest data). Appendix A.1 presents a detailed mathematical description of the algorithm, and a summary is given below:

- Compute a *k*-nearest-neighbor (*k*-NN) list for each data point.
- Compute a weighted *shared nearest-neighbor* (SNN) graph (Jarvis and Patrick, 1973) of the data based on the *k*-NN list. In the SNN graph, two data points are connected only if they belong to each others’ nearest-neighbor lists.
- From this graph, extract a high number of subgraphs by sparsification.
- Compute mean vectors of the connected components in each subgraph to obtain multiple sets of GMM mean-vector candidates.
- Choose a best set of initial mean vectors according to a minimum distance rule.

The method was validated against state-of-the-art-algorithms, such as Ward’s method (Ward, 1963), *K*-means (MacQueen, 1967), *K*-means++ (Arthur and Vassilvitskii, 2007) and Affinity propagation (Frey and Dueck, 2007). The validation results are presented in supplementary material (Section 3).

The proposed method depends on a single user parameter: a neighborhood size *k*. This parameter describes how many neighboring feature vectors (voxels) are used to form the SNN graphs.^5^ A choice of *k* affects the total number of clusters indirectly: Smaller values of *k* lead to large number of small clusters and thus can describe fine details of the original data. However, too detailed segmentation is difficult to grasp from the visualizations. Larger values of *k* lead to a lower number of clusters but to greater loss of the details of the data. Thus, a choice of *k* is a compromise between fine-graininess and interpretability of the findings. In this sense, *k* is not an ad hoc parameter but rather determines granularity level of the analysis.

We selected *k* as follows: First, we run FuSeISC for several values of *k*. Then, we plotted the total number clusters as a function of *k* and selected a value from the region where the number of clusters remained relatively constant (see Section 4 in supplementary material for validity of this approach using synthetic data). To confirm that the selected *k* value was appropriate, we also computed the similarity using the adjusted rand index (ARI) (Hubert and Arabie, 1985) between all FuSeISC solutions constructed from different values of *k* (see Section 2 in supplementary material for details of the ARI). In the constructed “stability matrix”, we looked for a stable region of high ARI values, because in this region the segmentation results were similar irrespective of the choice of *k*. Finally, we picked *k* from the region which showed stability in terms of both the total number of clusters and ARI.

### 3.3. Postprocessing

Large brain regions are not expected to be activated by the stimuli, which complicates the interpretation of the clustering results. An easy way to simplify the interpretation is to a priori discard voxels from the expected noise areas and run the FuSeISC only across voxels of interest^6^. This approach is also computationally much faster than the full-brain analysis. However, in this study we found it useful to evaluate the segmentation results across the whole brain for several reasons. First, the whole-brain segmentation serves as a validation tool for FuSeISC to detect large noise regions as separate clusters as well as to avoid mixing these areas with the activated cortical areas. Second, the distinction between “interesting” and “non-interesting” brain areas is not obvious: Although clusters located in the cerebral white-matter likely reflect noise, some interesting clusters may partially extend to these regions. Third, a whole-brain analysis tells whether our method can deal with large data in a feasible time (the StudyForrest data consisted of as many as 449,612 data points).

Because a whole-brain analysis leads to segmentation of both noise- and stimulus-related regions, we designed a postprocessing scheme to separate expected noise clusters from the clusters of interest. At first, we constructed a mask consisting of cerebral white-matter, brainstem, and ventricles, and counted how many voxels fell within this mask for each cluster. Then we sorted the clusters according to these counts and discarded clusters with highest counts from the rest of the analysis as noise. The exact number of discarded clusters was determined based on visual inspection of the spatial distributions of the clusters so that the clusters mainly distributed close or inside the noise mask were discarded. In the Results section, we concentrate on analyzing clusters of interest. The noise clusters are also briefly discussed and are displayed as supplementary material (Section 6).

We additionally discarded clusters reflecting border artifacts resulting from slightly different anatomical registration across the time series. These clusters were easily detected, because the mean ISC for at least one of the time series in these clusters was exactly zero.

### 3.4. Code availability

FuSeISC has been integrated to the ISC toolbox (Kauppi et al., 2014) and is freely available at https://www.nitrc.org/projects/isc-toolbox/.

## 4. Results

### 4.1. Comparison between conventional ISC and FuSeISC maps

It is insightful to compare FuSeISC maps with “conventional” univariate ISC maps. For this purpose, we computed conventional ISC maps across each five clip of interest for the StudyForrest data using the ISC toolbox (Kauppi et al., 2014). Thresholds for statistical significance were determined using a resampling procedure implemented in the toolbox. The thresholds were multiple comparison corrected across the voxels using the FDR (*q* < 0.001; the standard setting of the ISC toolbox).

Figure 2(A) shows three axial slices of the ISC map across Clip0. The colormaps denote ISCs averaged across all subject-pair-wise computations. The ISC is highest in the auditory cortex, which is expected because the stimuli were auditory. Interestingly, however, also frontal cortices show statistically significant ISC.

**Figure 2:**
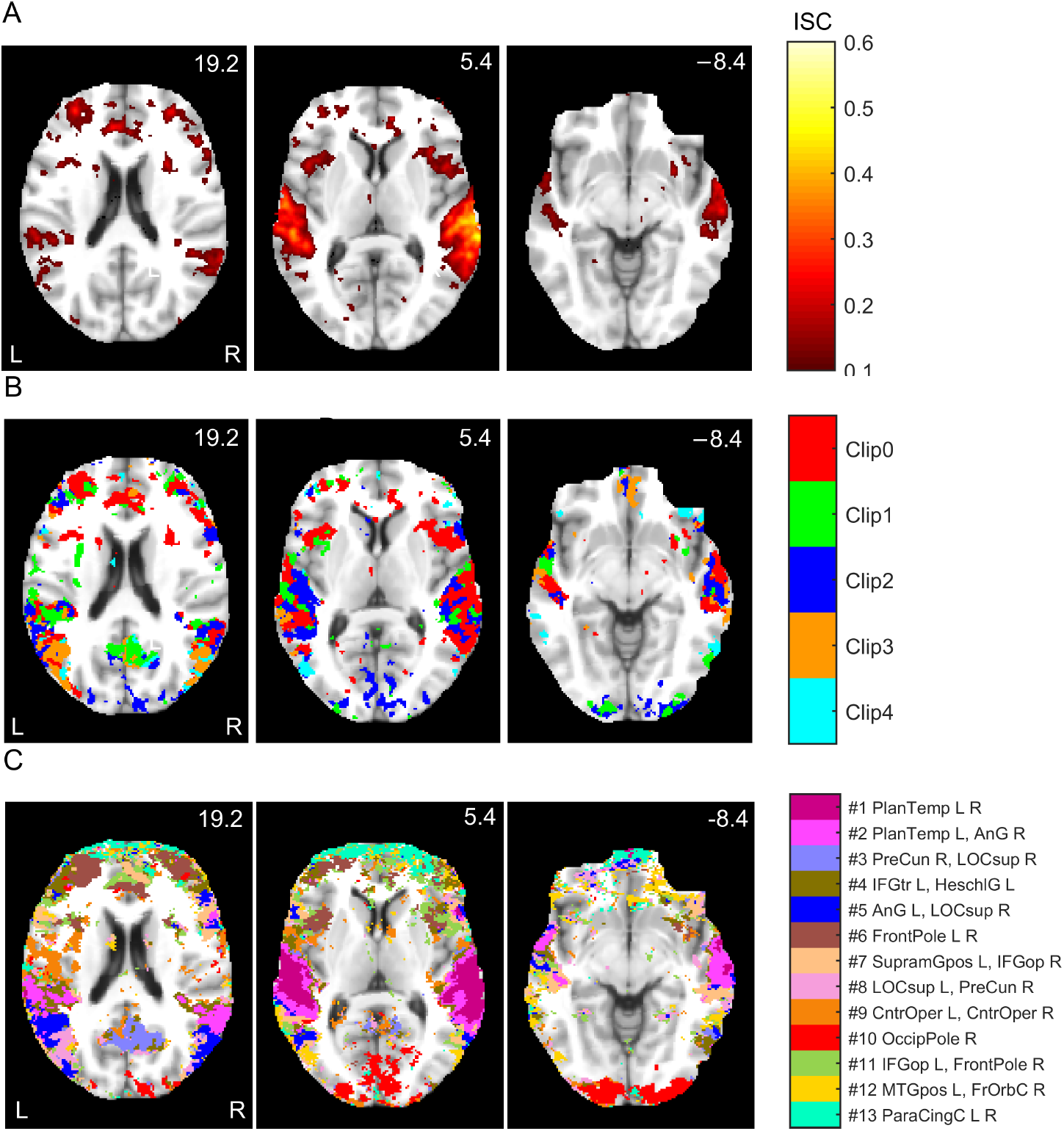
Comparison between conventional ISC and FuSeISC results for the StudyForrest data: (A) ISC map for Clip0, (B) Integrated ISC map of the five clips (when several clips resulted in statistically significant ISC in the same voxel, the voxel is color-coded according to the strongest ISC), and (C) FuSeISC map of the five clips. The axial slices are presented in millimeters in the MNI coordinates. The ISC maps were FDR corrected at *q* < 0.001 across all the voxels. FuSeISC does not require threshold selection, but clusters located dominantly over cerebral white-matter, brainstem, or ventricle areas were discarded. Note how FuSeISC found spatially meaningful segmentation and revealed more brain areas than conventional univariate ISC mapping.

Figure 2(B) superimposes the ISCs for all five clips. (Red color denotes statistically significant ISC during Clip0, green during Clip1, and so on; when several clips elicited significant ISC in the same voxel, the color code refers to the clip with the highest ISC.) All clips revealed statistically significant ISC in the auditory cortex, with right-hemisphere dominance, but the spatial location of ISCs also varied depending on the clip. For instance, Clip0 showed ISC in frontal regions whereas Clip2 showed ISCs in the posterior visual cortex (red and violet blobs, respectively).

Figure 2(C) shows a FuSeISC map of the same data using a neighborhood-size parameter *k* = 230 after postprocessing (see Section 4.4 on how we selected neighborhood sizes for the StudyForrest and ICBM data). Whereas conventional ISC mapping simply tells which voxels show statistically significant mean ISC across subject pairs for different clips, FuSeISC divides the brain into functional clusters formed on the basis of both mean and variability features of the subject-pair-wise ISCs extracted for each clip. Each cluster is shown in different color, and the names of the brain regions corresponding to the center of mass of the clusters are listed next to the colorbar. The names of the largest and the second largest subclusters are provided.^7^ For a more comprehensive listing of brain regions for each cluster, see supplementary material (Section 1, Table S3).

FuSeISC provided physiologically feasible functional division, with clusters in auditory and visual cortices. Many of the clusters were spatially local in one hemisphere and/or symmetric between the hemispheres, strongly suggesting that they reveal plausible brain processing instead of noise. Interestingly, FuSeISC revealed brain areas that remained undetected by the conventional ISC. For instance, some of the frontal regions covered by the FuSeISC map were not covered by the ISC maps of the individual clips in Fig. 2(B). Thus FuSeISC seemed to be more sensitive than the conventional ISC mapping for detecting activated brain areas.

### 4.2. Segmentation of StudyForrest data

To gain a better insight into the FuSeISC results, we divided the found clusters into two different spatial maps according to their relative ISC variability (i.e., ISC variability with respect to the ISC mean). The purpose of this division was to 1) highlight how low/high variability information is distributed across the brain, and 2) simply reduce the amount of information shown in a single brain image to make visual inspection of the results easier. We used the relative variability instead of the plain variability because the ISC variability was observed to increase with the mean, making the ranking of the clusters based on plain variability less interesting. The scatter plots of the mean and variability features are available in Fig. S8 (Section 7 in Supplementary material), confirming that the ISC variability increases with the mean. Increasing response variability together with a response mean has been previously reported in both animal and human brain signals (Tolhurst et al., 1981, Tanskanen et al., 2007). However, the increased variability of the ISC with increasing mean cannot be explained by general properties of the correlation coefficient because the variance of correlation coefficient decreases with increasing correlation (Bowley, 1928). This issue therefore deserves more thorough investigation in the future.

Clusters with low relative variability are expected to be found in early sensory areas where the processing is most coherent across subjects. Instead, clusters with high relative ISC variability are expected to be found both in the less coherent sensory areas and in higher-order brain areas that are involved in stimulus-related processing in a subject-dependent manner. The relative variability was computed as a fraction of the GMM mean vector elements for each cluster *i* as follows:

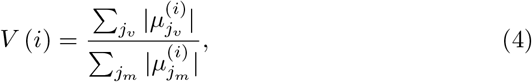

where *j*_*m*_ are the indexes of the ISC mean and *j*_*v*_ are the indexes of the ISC variability features in the model.

Fig. 3(A) shows spatial maps of the clusters with low relative ISC variability. These clusters were predominantly located in temporal lobes, especially covering the supratemporal auditory cortex. The supratemporal cluster was separated from the larger perisylvian cluster, as well as from a cluster in the temporoparietal junction. Fig. 3(B) shows clusters with high relative ISC variability. Most of the these clusters were located in frontal and occipital regions. The complete 3D spatial maps of clustering results are available in the NeuroVault service (Gorgolewski et al., 2015) at http://www.neurovault.org/collections//PXNGFJTL/.

**Figure 3:**
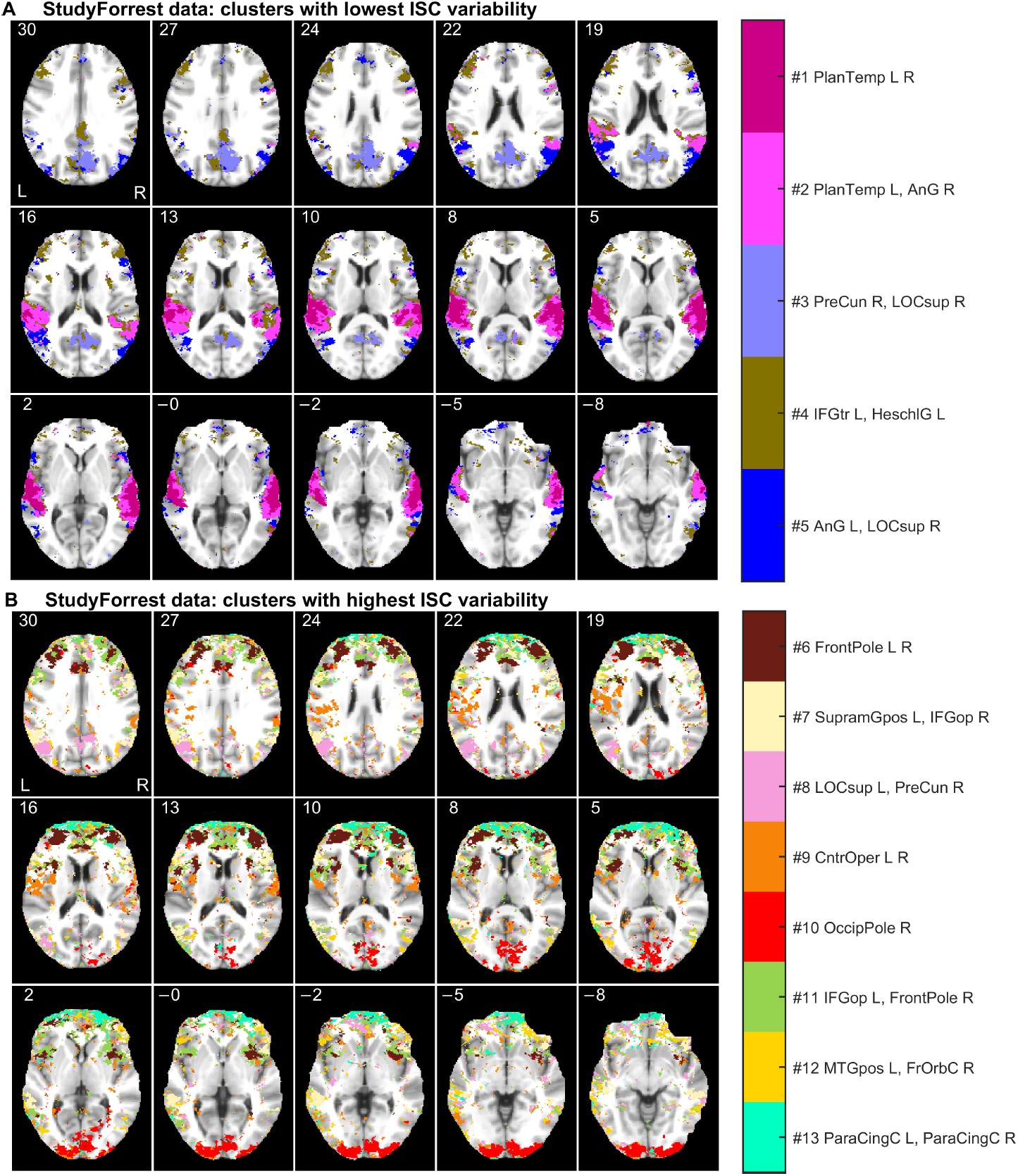
Functional segmentation of the StudyForrest data: (A) clusters with lowest ISC variability, and (B) clusters with highest ISC variability relative to the mean. The axial slices are presented and labeled with millimeters in the MNI coordinates. For the abbreviations of the brain region names and the spatial coordinates of the cluster centers, see supplementary material (Section 1, Tables S1 and S2).

In addition to spatial information, FuSeISC provides a characteristic pattern of ISC features (mean and variability) for each cluster, showing how the different stimulus sequences have contributed to each cluster. Figure 4 shows these “building blocks”, extracted from the estimated model of the StudyForrest data. Figures 4(A) and (B) correspond to the clusters of lowest and highest ISC variability (see Fig. 3), respectively. The contributions of the five audio clips on each cluster are coded in grayscale. For instance, temporal-lobe clusters showed highest ISCs during Clips 0–3 (see the first and the second bars of the mean ISC in Fig. 4(A)).

**Figure 4:**
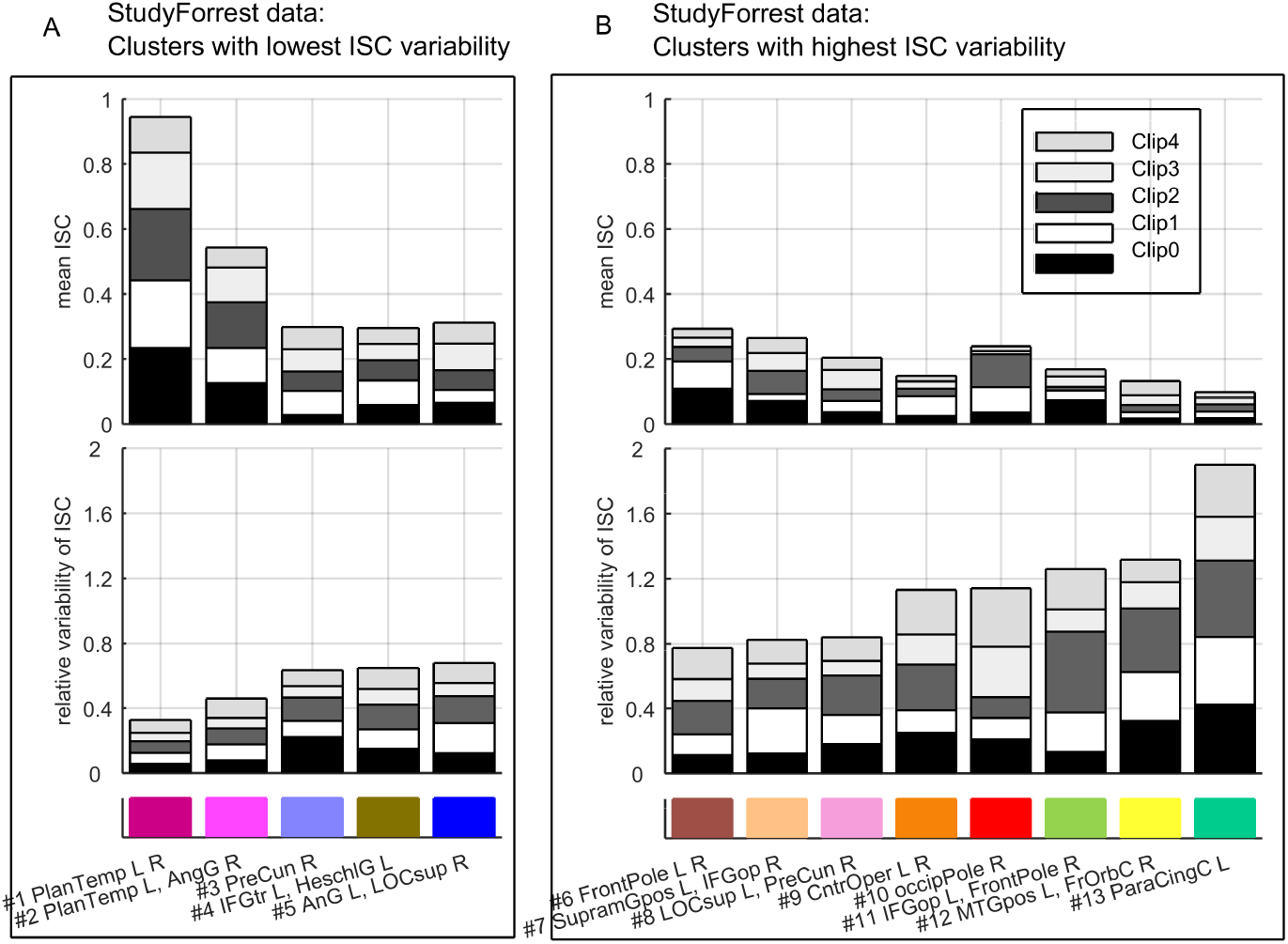
“Building blocks” of the clusters found in the StudyForrest data: (A) ISC features of clusters with lowest variability, and (B) ISC features of clusters with highest variability. Note that the clusters are ordered according to the *total* (relative) variability across the clips, meaning that the heights of the variability bars are in the increasing order. The feature values shown in the bars correspond to the distribution (mean) parameters of the GMM. The grayscale corresponds to audio clips of interest, and the color code corresponds to clusters shown in Fig. 3. See Table S1 in supplementary material for the abbreviations of the brain region names.

### 4.3. Segmentation of ICBM data

Fig. 5 shows spatial maps for the ICBM data (ICBM37ORIG). Similar to the StudyForrest data, clusters with low and high relative ISC variability are visualized separately. Clusters with low ISC variability were mainly located in the occipital lobes (see Fig. 5(A)), with segmentation of the visual cortices into multiple areas. This division of brain areas resembles results of independent component analysis (ICA) of fMRI data obtained during natural viewing (Pamilo et al., 2012), with different segments for foveal and peripheral vision, for example.

**Figure 5:**
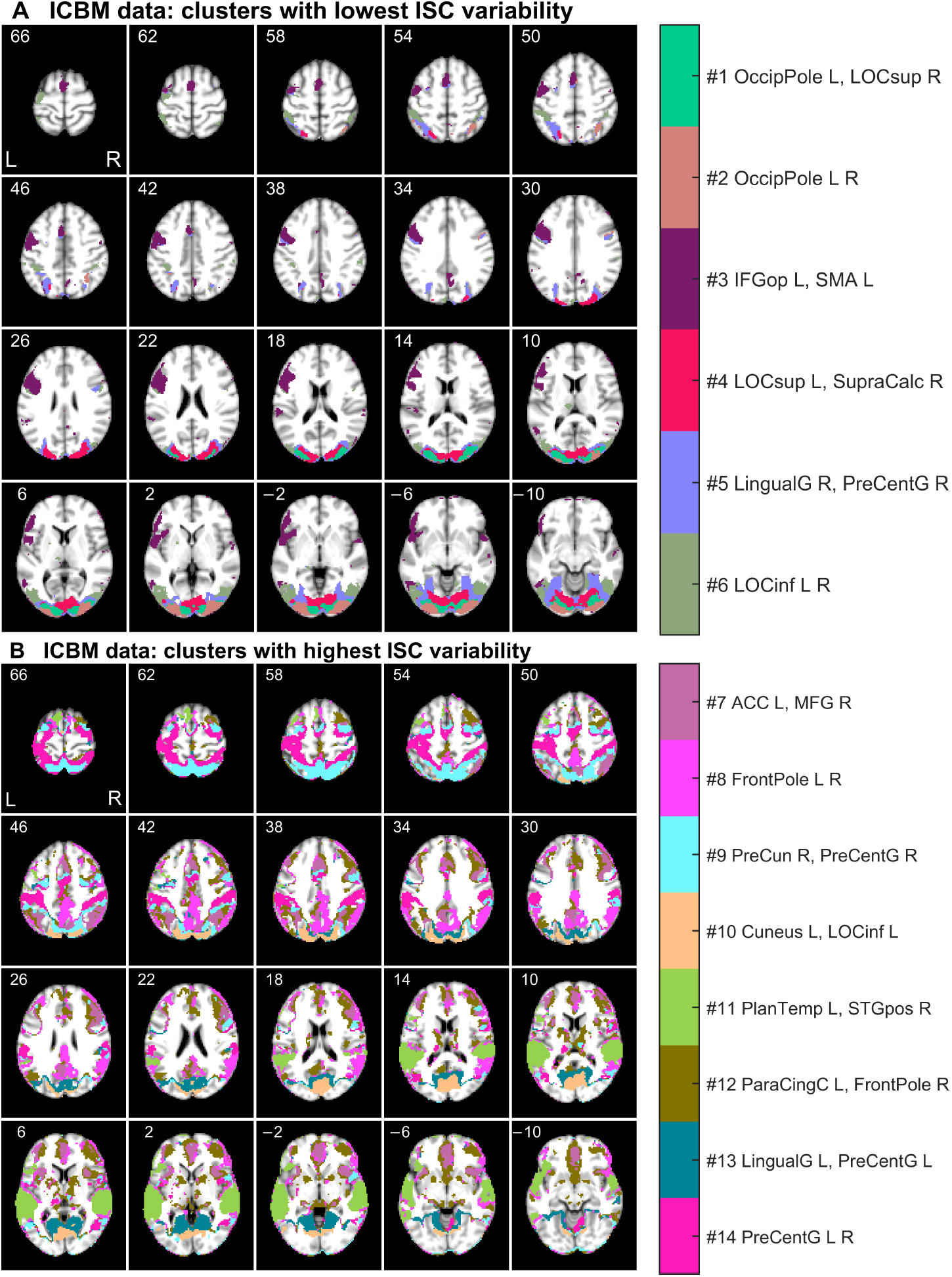
Functional segmentation of the ICBM data: (A) clusters with lowest ISC variability, and (B) clusters with highest ISC variability relative to the mean. The axial slices are presented and labeled with millimeters in the MNI coordinates. For the abbreviations of the brain region names and the spatial coordinates of the cluster centers, see supplementary material (Section 1, Tables S1 and S3).

Clusters with highest relative ISC variability were dispersed across many regions of the cortex (see Fig. 5(B)). For instance, separate clusters covered the intraparietal sulcus bilaterally, extrastriate body area, and parahippocampal space area. Interestingly, the segmentation also seemed to delineate nodes of the “default-mode network” in the posterior parietal cortex and medial prefrontal cortex).

Figure 6 shows the contributions of the five tasks on the ICBM clusters. Inspection of both the bar diagrams and the spatial locations of the clusters supports the physiological relevance of the obtained functional segmentation. For instance, clusters #1 and #2 in the visual cortices showed high mean ISC during external ordering (EO), hand imitation (HA), oculomotor (OM), and verb generation (VG) tasks. This result is unsurprising because these tasks were based on visual stimuli. In contrast, the large cluster #11 in the temporal lobe showed high mean ISC during the auditory naming (AN) task. Also this is physiologically plausible, because AN was the only task in which the stimuli were presented auditorily. Moreover, a cluster exhibiting high mean ISC during the hand imitation task was located around the sensorimotor strip (cluster #14).

**Figure 6:**
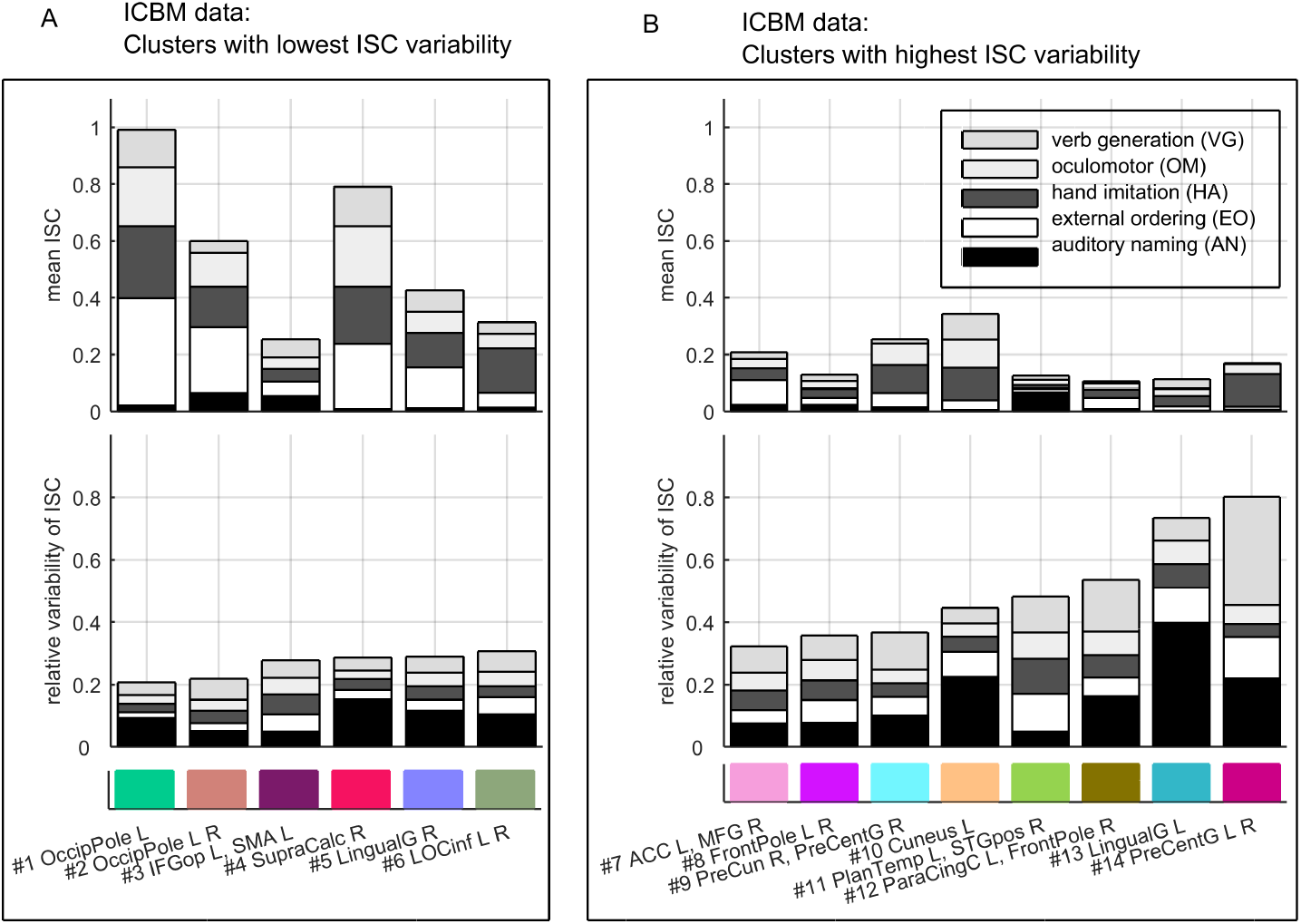
“Building blocks” of the clusters found in the ICBM data: (A) ISC features of clusters with lowest variability, and (B) ISC features of clusters with highest variability. Note that the clusters are ordered according to the *total* (relative) variability across the tasks, meaning that the heights of the variability bars are in the increasing order. The feature values shown in the bars correspond to the distribution (mean) parameters of the GMM. The grayscale corresponds to tasks of interest, and the color code corresponds to clusters shown in Fig. 5. See Table S1 in supplementary material for the abbreviations of the brain region names.

### 4.4. Selection of final segmentations

Here we describe how we selected *k* to obtain the final FuSeISC maps shown in the previous section. First, we ran FuSeISC for several values of *k* and plotted the total number of clusters for each result. Then, we found the range of stable values of *k* leading to a constant number of clusters. Fig. 7(A) shows the total number of clusters found for real ICBM and StudyForrest data sets as a function of a neighborhood size *k*. Interestingly, the curves were highly similar to each other. With small *k*-values, the number of clusters was high but the number decreases rapidly as *k* became larger. When *k ≥* 230, the number of clusters in the ICBM data stabilized around 20. For the StudyForrest data, the number of clusters in a stable region was approximately the same.

**Figure 7:**
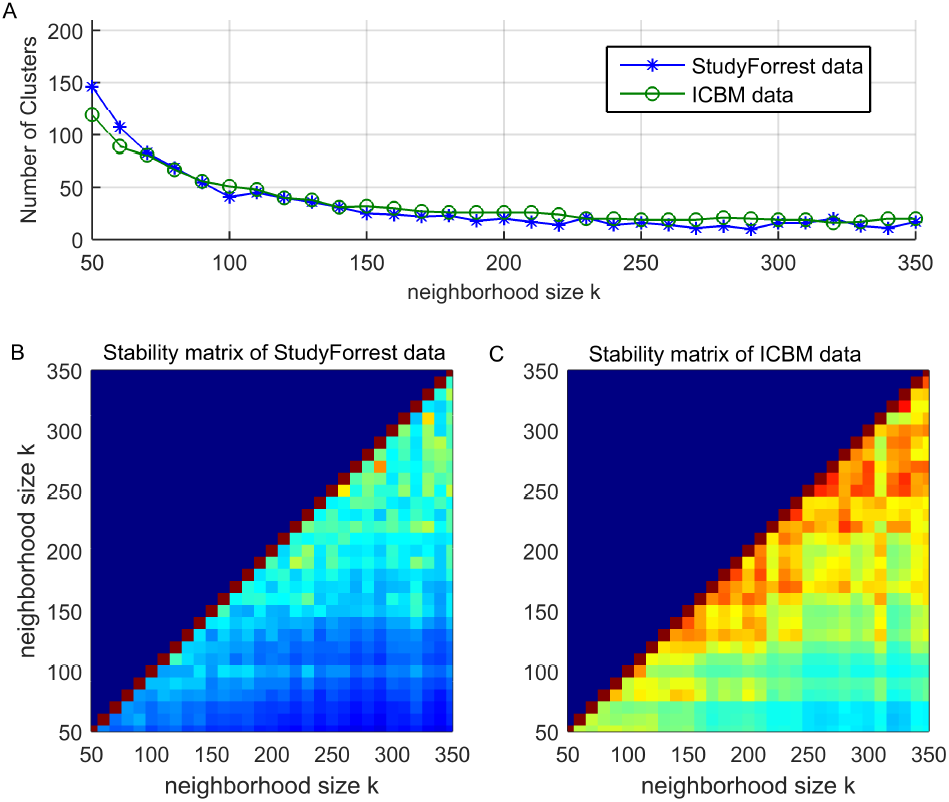
The effect of neighborhood size for the clustering results of the fMRI data: A) Total number of clusters of the ICBM and StudyForrest data, B) ARI stability matrix of the StudyForrest data, and C) ARI stability matrix of the ICBM data.

In addition, we computed ARI between results obtained for different values of *k*. In the resulting stability matrix, a high ARI value indicates that the segmentation result is stable, i.e., similar for two different choices of *k*. Figures 7(B) and (C) show the ARI stability matrices for the StudyForrest data and ICBM data, respectively. For both data sets, clustering solutions started to stabilize when *k* was relatively large (red color in the stability matrix indicates high similarity between the results computed for different values of *k*). The ARI values of the StudyForrest data were slightly lower than those of the ICBM data, which is expected because the spatial resolution (and the total number of voxels) in the StudyForrest data was notably higher.

Based on the above findings, we selected one of the stable solutions from both data sets for closer inspection (*k* = 250 for the ICBM data and *k* = 230 for the StudyForrest data). In these solutions, the exact number of clusters was 19 for the ICBM data and 21 for the StudyForrest data. After postprocessing, the total number of clusters was 14 for the ICBM data and 13 for the StudyForrest data.

#### Simulated ICBM data

To further validate our approach, we analyzed the simulated ICBM data and compared the results with the ground truth. Figure 8(A) presents the performance of the functional segmentation for the simulated ICBM data against the ground truth as a function of the neighborhood size *k*. For a wide range of parameters, ARI values resulted in “moderate agreement” (ARI between 0.4–0.6) between the ground truth and the estimated cluster labeling computed across the 72,577 voxels that were activated in the ground truth for at least one task. However, to make the clustering task realistic, FuSeISC was run across the entire brain involving 449,612 voxels.

**Figure 8:**
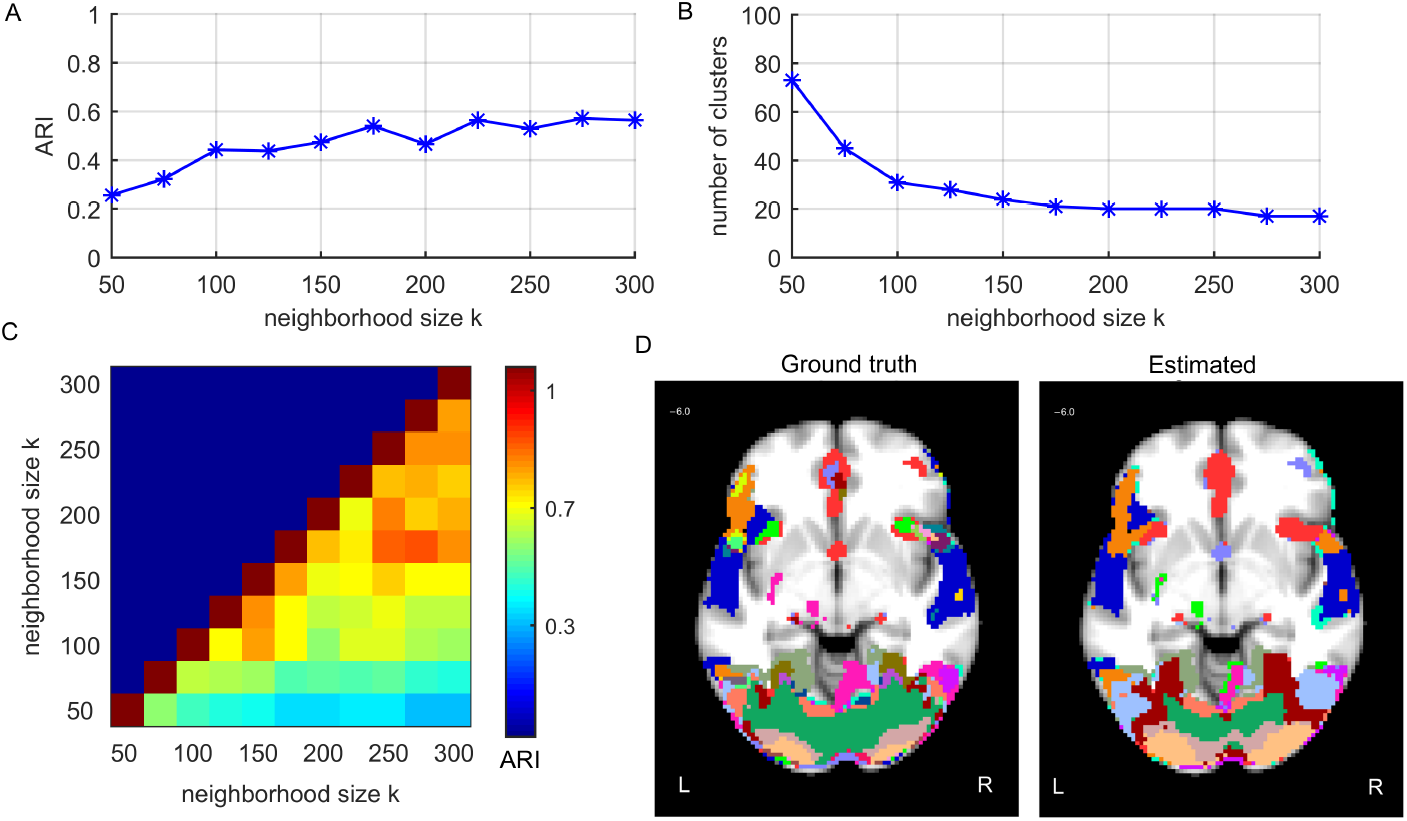
Results of the FuSeISC for the simulated ICBM data: A) clustering quality, B) total number of clusters, C) stability of the results, and D) an example slice showing spatial organization of the clusters (both ground truth and estimated clusters are shown).

Figure 8(B) shows the total number of clusters as a function of *k*. Clearly, the curve shows a region of constant number of clusters (20) when 200 ≤ *k* ≤ 250. This result corresponded well with the real-data results (see Fig. 7B), where the stable regions also consisted of about 20 clusters.

The ARI stability matrix of the solutions in Figure 8(C) shows the similarity between segmentation results computed for different values of *k*. Clearly, segmentation results were stable within the aforementioned constant region, since the ARI values were high.

Figure 8(D) shows a spatial organization of the clusters for one stable result (*k* = 225) and one axial slice (*z* = 6.0 mm in MNI). The ground-truth segmentation (left) and the estimated segmentation (right) are shown side by side to allow comparison. Based on visual inspection, the estimated segmentation resembles the true segmentation in most regions very well.

#### Results for ICBM data sets composed of different subjects

We also run FuSeISC for the two ICBM data sets consisting of different subjects (ICBM37#1 and ICBM37#2) and compared the obtained segmentations. Fig. 9 shows the spatial maps of both segmentations side-by-side; we show postprocessed segmentations, where white-matter and CSF clusters are eliminated, to simplify comparison. The corresponding raw segmentation results are available in Fig. S9 (Section 8 in supplementary material). To compare clusters between the data sets, we computed the Dice index (Dice, 1945) values between all the clusters in the two data sets (see Section 2 in supplementary material for details of the Dice index) and then used a Munkres assignment algorithm (Munkres, 1957) to match the clusters with each other. The Dice index values between the clusters are shown next to a color bar (“NaN” means that the corresponding cluster is present only in the leftmost data set). In many brain areas, the segmentation was visually very similar across the two data sets. The Dice index values between the clusters varied between 0.2–0.8. Using the same categorization for the Dice index as in Pajula et al. (2012), this result indicates slight to substantial agreement between individual clusters. The ARI value computed across the whole brain (228,483 voxels) between the two segmentations was 0.30.

**Figure 9:**
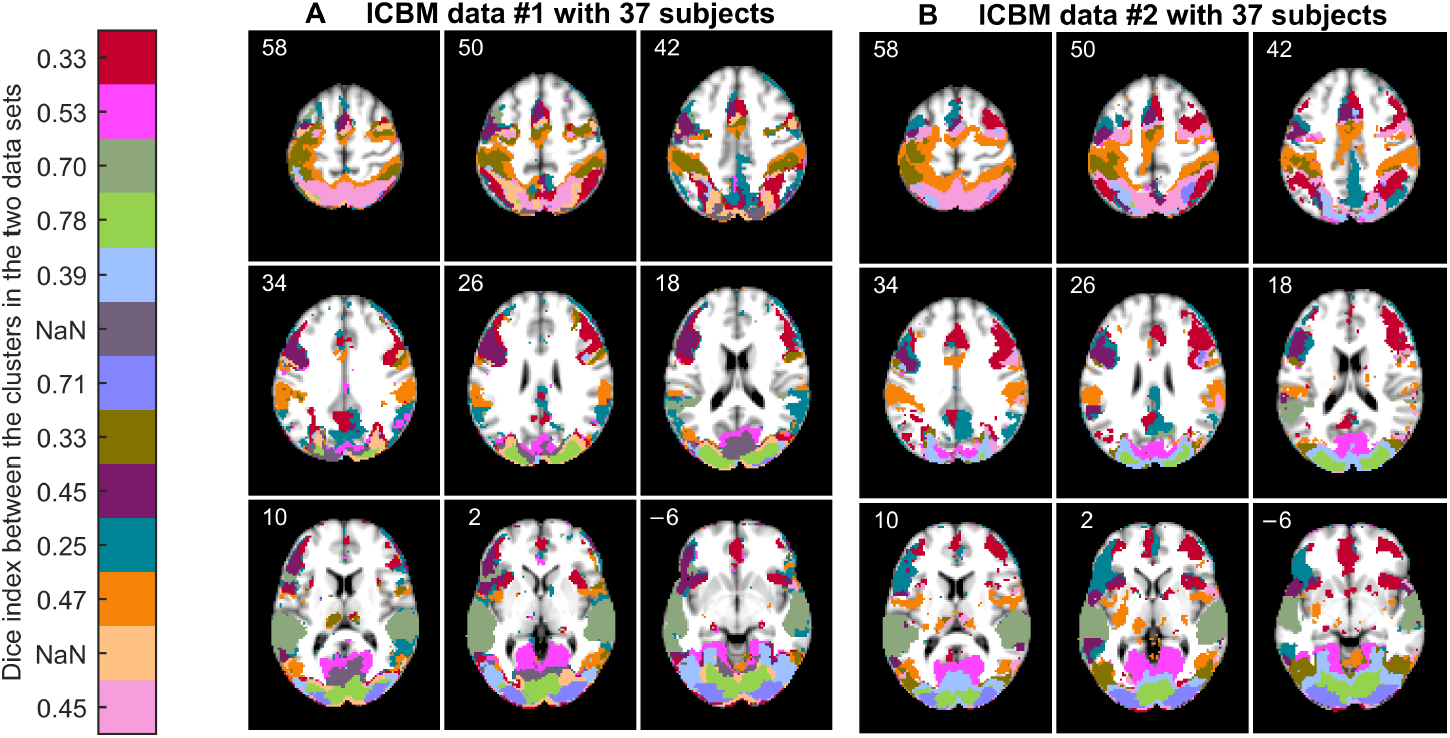
Functional segmentation result (*k* = 250) of (A) ICBM data set with 37 subjects, and (B) ICBM data set with another 37 subjects. The cortical segmentation in the two data sets is relatively similar. Clusters in the two data sets are matched using the Munkres assignment algorithm. Similarity between the clusters according to the Dice index is shown next to the color bar, “NaN” meaning that the corresponding cluster is present only in the leftmost data set. Postprocessing was used to emphasize similarities and differences between the results in the cortical areas. The corresponding raw segmentation results are provided in Fig. S9 (Section 8 in supplementary material).

Table 3 shows how the number of clusters in the FuSeISC results depends on the number of subjects used in the analysis. For both simulated and real ICBM data sets, the number of clusters increased together with the number of subjects. This result is plausible because the complexity of the data increases together with the number of subjects. For a fixed number of subjects, the number of clusters was quite similar across all three data sets. Slightly fewer clusters were found for the simulated data, as can be expected because we did not include ISC variability which is present in the real data. Table 3 also shows that the ARI values between the data sets containing different numbers of subjects varied between 0.27–0.52. For more detailed comparison between the clustering results with different number of subjects, see the spatial maps and cluster-wise similarities for the real ICBM data#2 in Figs. S10–S12 (see Section 8 in supplementary material).

**Table 3:**
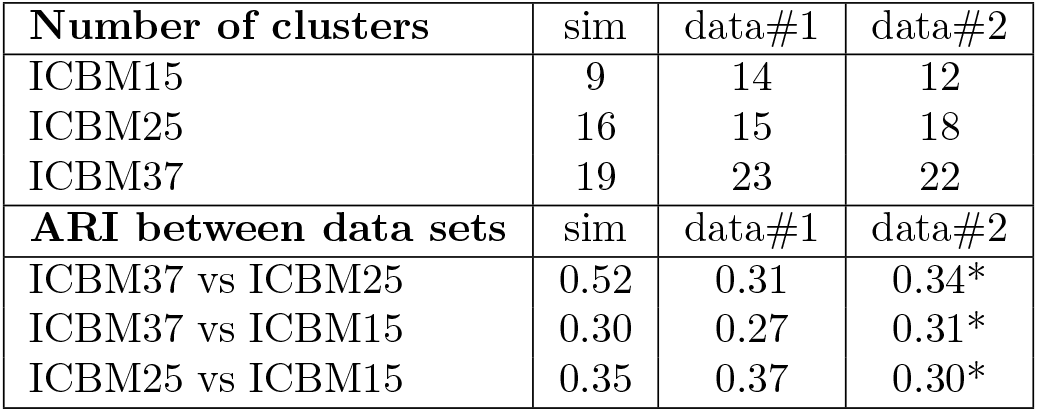
Effect of subject set size for the number of clusters found and ARI. Results for both simulated (sim) and real ICBM data sets (data#1 and data#2) are presented. ARI values for the real data sets were computed across the whole brain (228,483 voxels), and ARI values for the simulated data were computed across the activated brain areas (72,577 voxels). The neighborhood size used in the analysis was *k*=225 according to the previous simulation results. Spatial maps for more detailed comparison are available for the results marked with asterisks (Figs. S10–S12, see Section 8 in supplementary material).

| Number of clusters | sim | data#1 | data#2 |
| --- | --- | --- | --- |
| ICBM15 | 9 | 14 | 12 |
| ICBM25 | 16 | 15 | 18 |
| ICBM37 | 19 | 23 | 22 |
| ARI between data sets | sim | data#1 | data#2 |
| ICBM37 vs ICBM25 | 0.52 | 0.31 | 0.34* |
| ICBM37 vs ICBM15 | 0.30 | 0.27 | 0.31* |
| ICBM25 vs ICBM15 | 0.35 | 0.37 | 0.30* |

#### Comparison between ICBM and resting-state data

To validate that spatial structures found by FuSeISC result from stimulus-related brain activity, we also run FuSeISC with rfMRI data and compared the obtained segmentation with the ICBM data. Figure 10 shows segmentation results of both rfMRI (A) and the ICBM data (B). Here, we did not discard white-matter/CSF clusters as postprocessing to allow comparison of the segmentations across the whole brain. The segmentation results of the rfMRI data were very noisy whereas the segmentations of the ICBM data consisted of spatially connected and/or symmetric segments. ARI value between the ICBM and rfMRI data sets was 0.0, indicating disagreement between the segmentation results. Lack of spatial structure in the rfMRI suggests that connected/symmetric clusters found in the ICBM and StudyForrest data sets reflect similar stimulus-related brain activity across subjects rather than within-subject correlations which are present in the fMRI data even in the absence of external stimuli.

**Figure 10:**
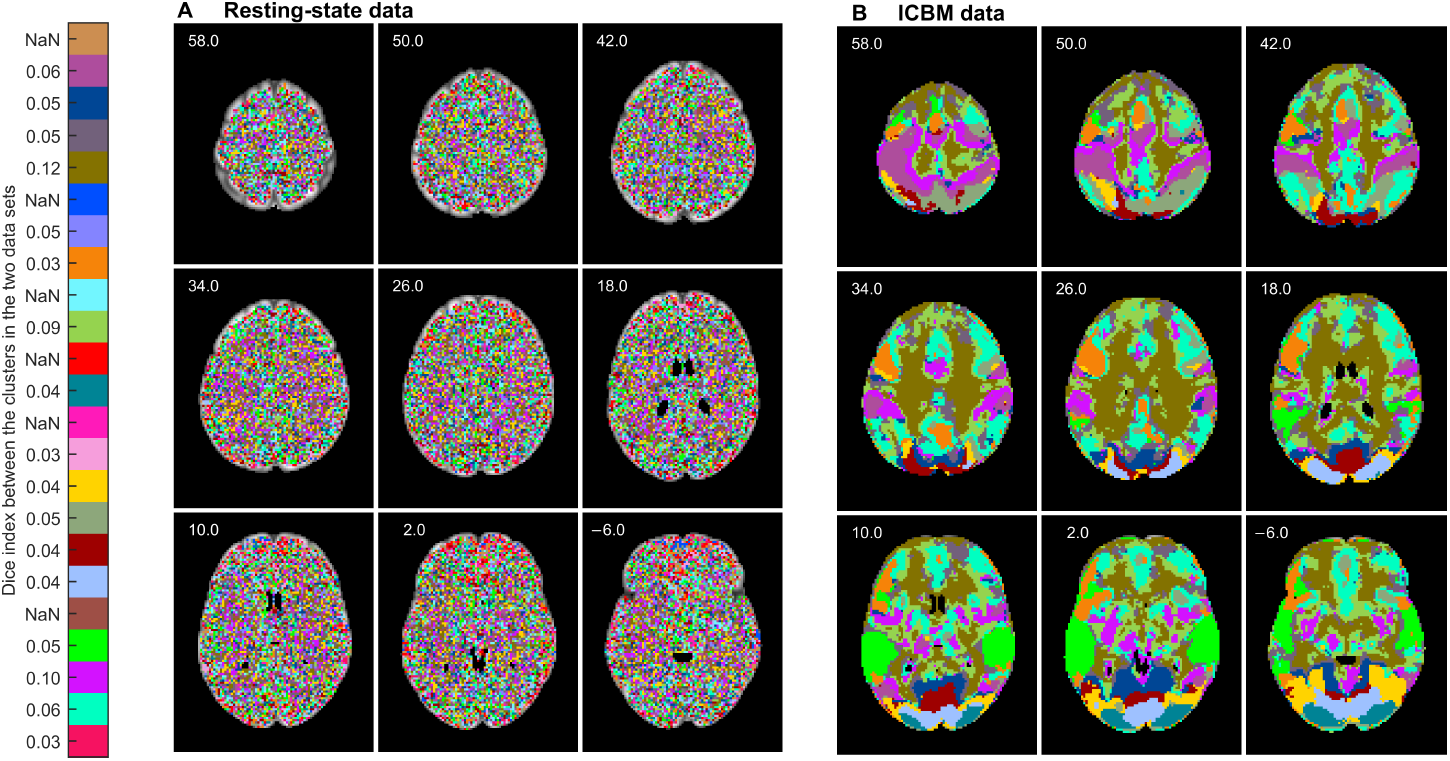
Functional segmentation result (*k* = 250) of (A) resting-state data, and (B) ICBM data. The segments of the resting-state are grainy whereas the segments of the ICBM data are spatially connected/symmetric. Clusters in the two data sets are matched using the Munkres assignment algorithm. Similarity between the clusters according to the dice index is shown next to the color bar, “NaN” meaning that the corresponding cluster is present only in the leftmost data set.

## 5. Discussion

### 5.1. Functional feasibility of the segmentation

The examination of the analysis results for the real fMRI data results in a couple of observations. Functional segmentations of both StudyForrest and ICBM data give an impression of a physiologically feasible division, with clusters in auditory, visual, and frontal cortices. Many of the clusters were symmetric between the hemispheres. Segmentations also seemed to delineate parts of the resting-state network. Although this network is considered to be highly “intrinsic” (Golland et al., 2007), it can be expected to change its state time-locked to the task demands and thereby show synchrony across subjects.

In FuSeISC, each cluster is characterized by its ISC mean and variability (bar plots in Figs. 4 and 6). In the ICBM data, the mean ISC patterns reflected well the expected brain areas involved in the tasks. Moreover, relative ISC variability of the clusters reflected, to at least some extent, the level of processing hierarchy in the brain. For instance in the StudyForrest data, clusters with high mean ISC but relatively low ISC variability were found in temporal lobes, where the majority of low-level auditory processing takes place during auditory stimuli. In contrast, clusters with higher relative ISC variability were found in higher-order brain areas, such as in the prefrontal areas and in the frontal poles. Based on visual inspection, the clusters were spatially relatively compact and/or bilaterally symmetric, indicating that also clusters with high ISC variability reflect real brain processing.^8^ This result is in line with recent studies indicating that the across-individuals variability in the functional brain areas and their connectivity carries meaningful information (Smith et al., 2014, Wang and Liu, 2014, Mueller et al., 2012, Gopal et al., 2016, Boldt et al., 2014, Zilles and Amunts, 2013).

The clusters for the 5-task ICBM data set covered most of the convexial and mesial cortices, thereby clearly extending the typical ISC maps that tend to concentrate on early sensory processing areas where the inter-subject correlations of fMRI time series are strongest because the activations are driven by the low-level sensory features of the stimuli (Kauppi et al., 2010a). These findings are in line with group-ICA results of fMRI obtained during natural viewing: the reconstruction of individual time courses shows considerably more inter-individual variability at, e.g., parieto-occipital sulcus than at early visual cortices (Malinen et al., 2007). The 5-task ICBM data set used in this work has been analyzed previously task-by-task for comparing the GLM and the conventional ISC (Pajula et al., 2012) for the purpose of validation of the ISC method. The analysis demonstrated that the active areas detected by the ISC (with no knowledge of the reference time course for the stimuli) and GLM (with a reference time course) were highly overlapping. In this work, all the tasks were analyzed jointly and one may thus ask whether the results would differ from just a combination of task-wise analysis. The visually most apparent difference was that the FuSeISC allowed the segmentation of visual cortex into multiple areas, as described in Results above, whereas in the conventional ISC analysis all tasks including visual input (verb generation, oculomotor, hand imitation, and external ordering) activated a large part of the visual cortex, with minor differences between the tasks.

Inter-subject variability can arise from several sources, one of them being between-subjects anatomical misalignment. To circumvent such challenges, between-subject alignment methods based on functional responses have been recently proposed (see Dubois and Adolphs (2016) for a review). In particular, fMRI during movie viewing has been found efficient in achieving correspondence via either maximizing inter-subject correlation (Sabuncu et al., 2010) or deriving a common representational space between subjects (Haxby et al., 2011, Guntupalli et al., 2016). These methods effectively reduce inter-subject variability in the data, but they may also mix spatial misalignment and intrinsic functional variability (Dubois and Adolphs, 2016), and they are costly in terms of scanning time. However, in future, functional segmentation methods might be developed to separate different types of between-subject variability.

### 5.2. Methodological considerations

Many cluster analysis techniques have been previously proposed for the functional segmentation of the human brain on the basis of fMRI data (Goutte et al., 1999, Craddock et al., 2012, van den Heuvel et al., 2008, Maggioni et al., 2014, Bellec et al., 2010, Blumensath et al., 2013, Eickhoff et al., 2016), but they have certain limitations in the analysis of complex group-fMRI data collected under diverse stimulation. Our method was particularly designed to address some of the key problems. For instance, conventional functional segmentation methods construct group-level segmentations by averaging results across individuals, ignoring inherent variability of brain functions across them. Importantly, FuSeISC does not cluster time series of the subjects themselves, but it computes and utilizes statistical information of the ISC features in clustering and this way naturally accounts both for similarity and variability in hemodynamic responses across subjects.

Previously, a clustering framework based on a two-layer generative model was introduced to account for inter-subject variability (Lashkari et al., 2012, Thirion et al., 2014). Unlike our cluster model built on the ISC features, that model utilizes information from the experimental setup. Wang et al. (2015) constructed functional segmentations separately for individuals using an iterative algorithm starting from the solution of the population atlas. While this approach takes into account individual differences, visual inspection of individual brain maps is a tedious task. One benefit of FuSeISC is that integrates data across all subjects and time series of interest into a single brain map and this way summarizes heterogeneous data into a meaningful amount of information for visual inspection.

Many existing functional segmentation methods constrain segmentation into spatially local neighborhoods (see e.g. Blumensath et al. (2013), Craddock et al. (2012)). FuSeISC does not assume that the clusters are spatially connected, but voxels are clustered without information about their spatial locations. This approach is plausible from neuroscientific perspective, as it allows to detect spatially distributed clusters as well as clusters with strikingly different sizes. Moreover, since spatial information is not used in the clustering process itself, visual inspection of the spatial locations of the clusters as well as spatial compactness of the clusters serves as a useful validation of the clustering outcome. The spatially compact clusters in our analyses indicate that the obtained segmentations reflect inherent structures of the fMRI data sets and not noise.

FuSeISC contains a user-definable parameter *k* which controls the coarseness of segmentation. Technically, *k* is used to decide the number of neighbors in *k*-NN lists and the subsequent optimally sparsified SNN graph. The graph, in turn, was used as a basis to initialize the GMM to improve the estimation accuracy. This way, *k* is only indirectly related the number of clusters that the clustering algorithm produces. Based on our simulations with synthetic Gaussian data, the value of *k* can be approximately interpreted as the number of voxels that each cluster should minimally contain (see Section 4 in Supplementary material). However, due to the complexity of real fMRI data, we proposed a systematic way to choose *k* based on the stability analysis of the number of clusters and similarity of the segmentation solutions (see Figs. 7 and 8(B–C)). For the tested data sets, the number of clusters stabilized close to 20 (irrespective of the data set), which is in line with Wang et al. (2015). On the other hand, it is possible that the number of found clusters varies notably between some data sets and the number of time series chosen, as the choice of optimal *k* depends on the intrinsic properties (size, shape, density and overlap of the clusters) of the data. In the future, we aim to study FuSeISC with a higher number of data sets and a different number of time series.

Smaller *k* values would result in more functional segments as illustrated in Fig. 7(A). Thus, for more detailed parcellations, a smaller *k* could be used. The smaller *k* values can be useful also to investigate some dedicated region of interest, either defined based on neuroanatomy or on a more coarse functional segmentation.

Due to the complex structure of the fMRI data, it is difficult to build an appropriate functional segmentation model in a general case. To alleviate the particular problems associated with the learning of the cluster model and selection of the total number of clusters, we proposed a new method based on SNN graph construction to initialize the GMM (see Appendix A.1). The method was successfully validated against the well-known methods *K*-means (MacQueen, 1967), *K*-means++ (Arthur and Vassilvitskii, 2007), Farthest first traversal algorithm (Hochbaum and Shmoys, 1985, Gonzalez, 1985), Ward’s minimum variance method (Ward, 1963), and Affinity propagation (Frey and Dueck, 2007) as well as its sparse version using simulated data sets containing Gaussian clusters and outliers (see Section 3 in supplementary material). These techniques were selected as they have been previously reported as useful in the initialization of the GMM, see for instance (Dueck, 2009, Dasgupta and Schulman, 2000, Fraley and Raftery, 2002, Blömer and Bujna, 2013). Moreover, all these methods can be conveniently controlled with a single user parameter, making them well-comparable against the proposed method. Although derived from a different point of view, we found very close correspondence in the clustering quality between our method and the affinity propagation algorithm. This finding was surprising and deserves further investigation. In any case, the benefit of our method over affinity propagation and Ward’s minimum variance method is that the full distance matrix needs not to be saved in the memory (even when spatial constraints are not used), allowing a large-scale segmentation across the whole brain without using spatial constraints.

FuSeISC is applicable for the analysis of large whole-brain multi-subject fMRI data sets as we have demonstrated in this paper. With a high-end desktop computer, the computation of the segmentation for one choice of *k* is feasible within 1–3 hours. However, we recommend optimizing the choice of *k* by running the FuSeISC segmentation for several values of *k*, possibly in parallel. When discarding a priori the voxels of the white-matter areas, brainstem and ventricles, computation time drops considerably. In this case, it is possible to evaluate several *k* values within few hours even without parallel processing.

The feature extraction step (involving computation of voxel-wise ISC matrices between each subject and estimation of the Jackknife ISC mean and variability estimates) is straightforward to parallelize for each time series. Note that when ISCs are computed between subjects using the ISC toolbox, there is no need to estimate threshold values for the ISC statistic using a block-bootstrap test, which is the computationally heaviest step in the conventional ISC analysis. We have added more details about computational demands of the initialization algorithm in the Appendix (see last section: Computational considerations).

### 5.3. Applications

In addition to being a tool for the spatial exploration of large fMRI data sets obtained using naturalistic stimulation (such as movies), FuSeISC has other potential applications. For example, it could be used to generate a functional atlas, either for a certain region of interest or for the whole brain, based on task-related fMRI.

This approach would be rather different than constructing atlases based on resting-state fMRI (see Craddock et al. (2012) and references therein) as, for example, fMRI-based functional connectivity patterns markedly depend on the brain state (Geerligs et al., 2015). As can be seen in Figs. 3 and 5, to achieve a resolution level of the currently commonly used resting-state fMRI atlases, a whole-brain atlas would require larger and more diverse data sets than the ones applied in this work. However, combined with a high-resolution fMRI of naturalistic experiments, our approach represents a novel line for future research. In principle, FuSeISC is not sensitive to the type of stimulus presentation, meaning that block-design, event-related, and naturalistic experiments could be combined together (at least when fMRI of the same set of subjects is acquired using the same scanner), partly facilitating atlas construction. Future research should show to what extent data combination is practically feasible.

As demonstrated in Figs. 4 and 6, FuSeISC also provides specific information about the ISC statistics of the time series of interest for each cluster, which can be used to trace clusters back to stimulus features. This is potentially useful if, for example, the multiple time series that form the input to FuSeISc are recorded during different scenes of a movie. Rich annotations of the stimulus sequence can then be used to relate clusters to different characteristics of the stimulus, providing an additional vehicle to interpret the FuSeISC parcellation.

FuSeISC also allows for reverse analysis, i.e., going back from the found clusters to the original ISC correlation matrices. The structures of the correlation matrices provide more details how the brains of different subjects have processed the same stimuli. For instance, high ISC variability may reflect subgroups of subjects who have dissimilar processing. Associating behavioral or other non-brain data with these subgroups or correlation matrices themselves (using, e.g., the Mantel test; see Salmi et al. (2013), Jääskeläinen et al. (2016)) could provide further insights into brain functions of different individuals.

Although FuSeISC is geared towards brain-imaging studies applying naturalistic stimulation, it can be equally well applied to traditional fMRI studies where the stimuli are strictly controlled. In the latter type of experiments, the results have been similar with ISC and standard GLM analyses (Pajula et al., 2012, Pajula and Tohka, 2014). However, it should be noted that the ISC method requires that the subjects received identical stimuli and therefore, FuSeISC is not useful in segmenting resting-state fMRI data that can be analyzed for example by using group-ICA (Beckmann et al., 2005, Kiviniemi et al., 2009).

## 6. Conclusions

We have proposed a new data-driven method, functional brain segmentation using inter-subject correlation, FuSeISC, to analyze fMRI data sets collected from a group of subjects who experience a variety of stimuli. The method segregates brain areas based on the ISC information without explicit knowledge of the stimuli. This way, FuSeISC clusters brain areas directly on the basis of a single data set formed from a group of subjects. Each cluster is characterized by its spatial location as well as by its specific ISC mean and variability. These properties make FuSeISC rather different from conventional functional segmentation algorithms and ISC analysis methods designed for fMRI data. The method is not only prominent for spatial exploration of large fMRI data sets obtained using naturalistic stimuli, but has also other potential applications such as generation of a functional brain atlases including both lower-and higher-order processing areas.

## Appendix A. Construction of initial Gaussian mixture model

## Appendix A.1. Generation of candidate models

Here we describe a simple but efficient technique for restricting a set of initial Gaussian mixture model (GMM) candidates *a priori*. To find good candidate models, we capture intrinsic structure of the data by *shared nearest-neighbor* (SNN) graphs (Jarvis and Patrick, 1973) (also called mutual nearest-neighbor graphs). In the SNN graph, two data points are connected only if they belong to each other’s *k*-nearest-neighbor sets. More formally, let us denote the set of *L* data points in a *d*-dimensional feature space as 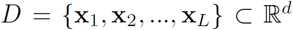, and let the set of the *k*-nearest neighbors^9^ of an arbitrary data point **x**_*m*_ be *N*_*m*_. In the SNN graph *G* (*D, E*), the vertex set *D* contains all the data points and the edge set *E* is given as follows (Jarvis and Patrick, 1973):

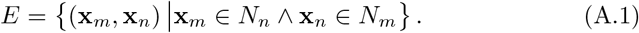

Furthermore, we weight every edge in *E* of the SNN graph by counting the total number of intersecting data points of the two nearest-neighbor sets:

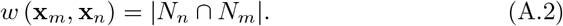

Note that by using this weighting scheme, the similarity between two connected data points does not depend on their *absolute distance* but the similarity between data points is determined by the *similarity of the k-nearest-neighbor sets of these data points*. This desirable property allows detection of clusters with varying densities even in a high-dimensional feature space (Tan et al., 2014, Ertöz et al., 2003, Houle et al., 2010). We also compute a *degree* for each data point **x**_*m*_ as the sum of the weights of edges connecting **x**_*m*_ and its nearest neighbors:

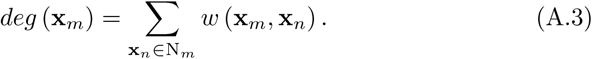

Next, we form multiple candidate (sub)graphs through *sparsification* of the weighted SNN graph. More specifically, to form a single candidate, we remove all the edges associated with data points **x**_*m*_ whose degree values are below a selected threshold *T*_*j*_. Several candidates are formed using multiple thresholds *T*_*j*_, for *j* = 1, 2*,…, q*.^10^ Thus, a final set of candidate graphs is:

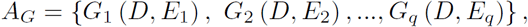

where the edge sets of the candidate graphs are:

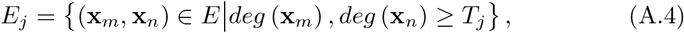
for *j* = 1, 2,*…, q*. Finally, we locate the centers of the connected components in each candidate graph:

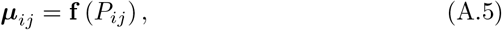
for *i* = 1, 2*,…, h*_*j*_. In this expression, ***µ***_*ij*_ denotes the center of the *i* th connected component in the *j*th graph *G*_*j*_, the set *P*_*ij*_ contains all the data points associated with that component, and *h*_*j*_ is the total number of connected components in that graph. The function **f** (·) defines a center of a connected component in a meaningful way. Our default choice for **f** (·) is the mean of the data points of the *P*_*ij*_.

## Appendix A.2. Choice of initial GMM

Given the candidate sets *C*_1_*, C*_2_*,…, C*_*q*_ of the mean vectors, the next task is to choose one set 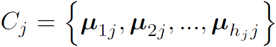 that represents all clusters in data. Different criteria can be used for this purpose, including well-known Bayesian information criterion (BIC) (Schwarz et al., 1978) or simple minimum sum-of-squared error (SSE) criterion (minimum distance rule). In our tests with synthetic noisy fMRI data, we found slightly more stable clustering results with the SSE than BIC (see Section 5 in supplementary material) and therefore we used SSE as the criterion in this paper.^11^

After selecting the best candidate set of mean vectors, we used the *K*-means algorithm (MacQueen, 1967) to update the mean vectors and assign all the data points to the clusters. Mixture weights were initialized by computing fractions of the data points within the clusters and covariance matrices were estimated from the data within the clusters. The obtained mean vectors, mixing weights and covariance matrices formed our initial GMM.

## Appendix A.3. Computational considerations

The construction of the *k*-NN graph in our initialization algorithm requires computation of *L*^2^ distances, where *L* is the number of data points. Memory requirement of the *k*-NN graph is O(*Lk*), which is not a problem since *k* ≪ *L* in practice. The computation time of the initialization algorithm is also dependent on the number of sparsification thresholds evaluated, which in turn depends on *k* and the properties of data. We have noted that for large data sets (which is the case with the fMRI data), the evaluation of all sparsification thresholds is computationally very heavy. Therefore, we use the following heuristic to analyze fMRI data: At first, we evaluate the SSE for every *k*th sparsification threshold. After this, we pick the best two SSE values, and evaluate all unique sparsification threshold values between the two thresholds to improve the SSE. To further save computation time, computationally most demanding steps (construction of the *k*-NN graph, selection of the best sparsified SNN graph) of our initialization algorithm are written in C language.

## Author Contributions

Conceived and designed the experiments: JPK, JP, JT. Performed the experiments: JPK, JP. Analyzed the data: JPK, RH, JT. Wrote the paper: JPK, JP, RH, JT. Contributed new clustering algorithm: JPK, JN.

## Acknowledgments

JPK was funded by the Academy of Finland Postdoctoral Researcher program (Research Council for Natural Sciences and Engineering; grant number 286019).

RH was funded by the Finnish Cultural Foundation (Eminentia grant).

JT received funding from the Universidad Carlos III de Madrid, the European Union’s Seventh Framework Programme for research, technological development and demonstration under grant agreement nr 600371, el Ministerio de Econom´ıa y Competitividad (COFUND2013-40258) and Banco Santander.

Data collection and sharing for this project were provided in part by the International Consortium for Brain Mapping (ICBM; Principal Investigator: John Mazziotta, M.D., Ph.D.). ICBM funding was provided by the National Institute of Biomedical Imaging and BioEngineering. ICBM data are disseminated by the Laboratory of Neuro Imaging at the University of Southern California.

The rfMRI data in this study were provided [in part] by the Human Connectome Project, WU-Minn Consortium (Principal Investigators: David Van Essen and Kamil Ugurbil; 1U54MH091657) funded by the 16 NIH Institutes and Centers that support the NIH Blueprint for Neuroscience Research; and by the McDonnell Center for Systems Neuroscience at Washington University.

### Conflicting interests

The authors declare no conflicting interests.

## Footnotes

1 This assumption is also made in model-based brain-mapping methods, such as those based on a general linear model (GLM; Friston et al. (1994))

2 http://studyforrest.org/pages/challenge.html, http://studyforrest.org/contest_fuseisc.html

3 http://www.loni.usc.edu/ICBM/Downloads/Downloads_FRB.shtml

4 This difficulty follows from the non-convexity of the maximum likelihood cost function to be minimized and every local optimization algorithm (including gradient methods) have this problem.

5 It is important to note that the connected components of the SNN graph are found in a *feature space* and not in a spatial domain and this way a single cluster may consist of multiple spatially connected components (subclusters).

6 This option is available in the ISC toolbox.

7 Because spatial constraints are not used in FuSeISC, each found cluster in a feature space can consist of more than one spatially disjoint subclusters. The name of the second largest subcluster is reported only when the actual cluster consists of at least two spatially disjoint subclusters whose sizes are greater than 100 voxels. Moreover, if the center of mass is located in white-matter or non-specified brain area, the largest cortical brain region intersecting with the cluster is reported instead of the location of the center of mass.

8 However, because of the higher spatial resolution of the fMRI recordings and sharper smoothing kernel (with smaller FWHM) during the analysis, some of the clusters were spatially more fragmented in StudyForrest than ICBM data.

9 A point is not its own neighbor, i.e. x^*m*^ ∉*N*^*m*^.

10 A most systematic approach is to construct as many candidates as there are distinct degree values. Note that degree values are integers and the maximum possible value is *k* (*k* − 1).

11 We found out that the SSE criterion returns meaningful solutions in which all clusters are represented by estimated centroids ***µ*** but these solutions did not trivially coincide with the solutions having highest number of clusters in the formed candidate sets. This can be explained by varying locations of estimated centroids between different candidate models.

## References

Abrams, D. A., Ryali, S., Chen, T., Chordia, P., Khouzam, A., Levitin, D. J., and Menon, V. (2013). Inter-subject synchronization of brain responses during natural music listening. Eur J Neurosci, 37(9):1458–1469.

Arthur, D. and Vassilvitskii, S. (2007). k-means++: the advantages of careful seeding. In Proc Symp Discrete Algorithms, pages 1027–1035.

Beckmann, C. F., DeLuca, M., Devlin, J. T., and Smith, S. M. (2005). Investigations into resting-state connectivity using independent component analysis. Philos Trans R Soc Lond B Biol Sci, 360(1457):1001–1013.

Bellec, P., Rosa-Neto, P., Lyttelton, O. C., Benali, H., and Evans, A. C. (2010). Multi-level bootstrap analysis of stable clusters in resting-state fMRI. NeuroImage, 51(3):1126–1139.

Blömer, J. and Bujna, K. (2013). Simple methods for initializing the EM algorithm for Gaussian mixture models. arXiv preprint arXiv:1312.5946.

Blumensath, T., Jbabdi, S., Glasser, M. F., Van Essen, D. C., Ugurbil, K., Behrens, T. E., and Smith, S. M. (2013). Spatially constrained hierarchical parcellation of the brain with resting-state fMRI. NeuroImage, 76:313–324.

Boldt, R., Seppä, M., Malinen, S., Tikka, P., Hari, R., and Carlson, S. (2014). Spatial variability of functional brain networks in early-blind and sighted subjects. NeuroImage, 95:208–216.

Bowley, A. (1928). The standard deviation of the correlation coefficient. J Amer Statist Assoc, 23(161):31–34.

Craddock, R. C., James, G., Holtzheimer, P. E., Hu, X. P., and Mayberg, H. S. (2012). A whole brain fMRI atlas generated via spatially constrained spectral clustering. Human Brain Mapp, 33(8):1914–1928.

Dasgupta, S. and Schulman, L. J. (2000). A two-round variant of EM for Gaussian mixtures. In Proc 16th Conf Uncertainty in Artificial Intelligence, pages 152–159.

Dice, L. R. (1945). Measures of the amount of ecologic association between species. Ecology, 26(3):297–302.

Dubois, J. and Adolphs, R. (2016). Building a science of individual differences from fMRI. Trends Cogn Sci, 20(6):425–443.

Dueck, D. (2009). Affinity Propagation: Clustering data by passing messages. PhD thesis, University of Toronto.

Eickhoff, S., Thirion, B., Varoquaux, G., and Bzdok, D. (2016). Connectivity-Based Parcellation: Critique and Implications. Hum Brain Mapp, page 22.

Ertöz, L., Steinbach, M., and Kumar, V. (2003). Finding clusters of different sizes, shapes, and densities in noisy, high dimensional data. In Proc 2nd SIAM Int Conf on Data Mining.

Essen, D. V., Ugurbil, K., Auerbach, E., Barch, D., Behrens, T., Bucholz, R., Chang, A., Chen, L., Corbetta, M., Curtiss, S., Penna, S. D., Feinberg, D., Glasser, M., Harel, N., Heath, A., Larson-Prior, L., Marcus, D., Michalareas, G., Moeller, S., Oostenveld, R., Petersen, S., Prior, F., Schlaggar, B., Smith, S., Snyder, A., Xu, J., and Yacoub, E. (2012). The Human Connectome Project: a data acquisition perspective. NeuroImage, 62(4):2222–2231.

Figueiredo, M. A. F. and Jain, A. K. (2002). Unsupervised learning of finite mixture models. IEEE Trans Pattern Anal Mach Intell, 24(3):381–396.

Fraley, C. and Raftery, A. E. (2002). Model-based clustering, discriminant analysis, and density estimation. J Am Stat Ass, 97(458):611–631.

Frey, B. and Dueck, D. (2007). Clustering by passing messages between data points. Science, 315(5814):972–976.

Friston, K. J., Holmes, A. P., Worsley, K. J., Poline, J.-P., Frith, C. D., and Frackowiak, R. S. (1994). Statistical parametric maps in functional imaging: a general linear approach. Hum Brain Mapp, 2(4):189–210.

Geerligs, L., Rubinov, M., Henson, R. N., et al. (2015). State and trait components of functional connectivity: individual differences vary with mental state. J Neurosci, 35(41):13949–13961.

Glasser, M. F., Sotiropoulos, S. N., Wilson, J. A., Coalson, T. S., Fischl, B., Andersson, J. L., Xu, J., Jbabdi, S., Webster, M., Polimeni, J. R., Essen, D. C. V., and Jenkinson, M. (2013). The minimal preprocessing pipelines for the Human Connectome Project. NeuroImage, 80:105–124.

Golland, Y., Bentin, S., Gelbard, H., Benjamini, Y., Heller, R., Nir, Y., Hasson, U., and Malach, R. (2007). Extrinsic and intrinsic systems in the posterior cortex of the human brain revealed during natural sensory stimulation. Cereb Cortex, 17(4):766–777.

Gonzalez, T. (1985). Clustering to minimize the maximum intercluster distance. Theor Comp Sci, 38:293–306.

Gopal, S., Miller, R. L., Baum, S. A., and Calhoun, V. D. (2016). Approaches to capture variance differences in rest fMRI networks in the spatial geometric features: Application to schizophrenia. Front Neurosci, 10.

Gorgolewski, K. J., Varoquaux, G., Rivera, G., Schwartz, Y., Sochat, V. V., Ghosh, S. S., Maumet, C., Nichols, T. E., Poline, J.-B., Yarkoni, T., Margulies, D. S., and Poldrack, R. A. (2015). Neurovault.org: A repository for sharing unthresholded statistical maps, parcellations, and atlases of the human brain. NeuroImage, 24:1242–1244.

Goutte, C., Toft, P., Rostrup, E., Nielsen, F. Å., and Hansen, L. K. (1999). On clustering fMRI time series. NeuroImage, 9(3):298–310.

Guntupalli, J. S., Hanke, M., Halchenko, Y. O., Connolly, A. C., Ramadge, P. J., and Haxby, J. V. (2016). A model of representational spaces in human cortex. Cereb Cortex, 26(6):2919–2934.

Hanke, M., Baumgartner, F. J., Ibe, P., Kaule, F. R., Pollmann, S., Speck, O., Zinke, W., and Stadler, J. (2014). A high-resolution 7-Tesla fMRI dataset from complex natural stimulation with an audio movie. Scientific Data, 1.

Hasson, U., Malach, R., and Heeger, D. J. (2010). Reliability of cortical activity during natural stimulation. Trends Cogn Sci, 14(1):40–48.

Hasson, U., Nir, Y., Levy, I., Fuhrmann, G., and Malach, R. (2004). Intersubject synchronization of cortical activity during natural vision. Science, 303(5664):1634–1640.

Haxby, J. V., Guntupalli, J. S., Connolly, A. C., Halchenko, Y. O., Conroy, B. R., Gobbini, M. I., Hanke, M., and Ramadge, P. J. (2011). A common, high-dimensional model of the representational space in human ventral temporal cortex. Neuron, 72(2):404–416.

Hochbaum, D. S. and Shmoys, D. B. (1985). A best possible heuristic for the k-center problem. Math Oper Res, 10(2):180–184.

Houle, M., Kriegel, H.-P., Kröger, P., Schubert, E., and Zimek, A. (2010). Can shared-neighbor distances defeat the curse of dimensionality? In Proc 22nd Conf on Scientific and Statistical Database Management (SSDBM), pages 482–500.

Hubert, L. and Arabie, P. (1985). Comparing partitions. J Classification, 2(1):193–218.

Jääskeläinen, I. P., Pajula, J., Tohka, J., Lee, H.-J., Kuo, W.-J., and Lin, F.-H. (2016). Brain hemodynamic activity during viewing and re-viewing of comedy movies explained by experienced humor. Sci Rep, 6.

Jarvis, R. A. and Patrick, E. A. (1973). Clustering using a similarity measure based on shared near neighbors. IEEE Trans Comput, 100(11):1025–1034.

Kauppi, J.-P., Jääskeläinen, I., Sams, M., and Tohka, J. (2010a). Clustering inter-subject correlation matrices in functional magnetic resonance imaging. In 10th IEEE Int Conf on Information Technology and Applications in Biomedicine (ITAB), pages 1–6.

Kauppi, J.-P., Jääskeläinen, I. P., Sams, M., and Tohka, J. (2010b). Intersubject correlation of brain hemodynamic responses during watching a movie: localization in space and frequency. Front Neuroinform, 4(0):5.

Kauppi, J.-P., Pajula, J., and Tohka, J. (2014). A Versatile software package for inter-subject correlation based analyses of fMRI. Front Neuroinform, 8(2).

Kiviniemi, V., Starck, T., Remes, J., Long, X., Nikkinen, J., Haapea, M., Veijola, J., Moilanen, I., Isohanni, M., Zang, Y.-F., et al. (2009). Functional segmentation of the brain cortex using high model order group PICA. Human Brain Mapp, 30(12):3865–3886.

Lashkari, D., Sridharan, R., Vul, E., Hsieh, P.-J., Kanwisher, N., and Golland, P. (2012). Search for patterns of functional specificity in the brain: a nonparametric hierarchical Bayesian model for group fMRI data. NeuroImage, 59(2):1348–1368.

MacQueen, J. B. (1967). Some methods of classification and analysis of multivariate observations. In Proc 5th Berkeley Symp Mathematical Statistics and Probability, pages 281–297.

Maggioni, E., Tana, M. G., Arrigoni, F., Zucca, C., and Bianchi, A. M. (2014). Constructing fMRI connectivity networks: a whole brain functional parcellation method for node definition. J Neurosci Method, 228:86–99.

Malinen, S., Hlushchuk, Y., and Hari, R. (2007). Towards natural stimulation in fMRI—issues of data analysis. NeuroImage, 35(1):131–139.

Marcus, D., Harwell, J., Olsen, T., Hodge, M., Glasser, M., Prior, F., Jenkinson, M., Laumann, T., Curtiss, S., and Van Essen, D. (2011). Informatics and data mining tools and strategies for the Human Connectome Project. Front Neuroinform, 5:4.

Mazziotta, J., Toga, A., Evans, A., Fox, P., Lancaster, J., Zilles, K., Woods, R., Paus, T., Simpson, G., Pike, B., Holmes, C., Collins, L., Thompson, P., MacDonald, D., Iacoboni, M., Schormann, T., Amunts, K., Palomero-Gallagher, N., Geyer, S., Parsons, L., Narr, K., Kabani, N., Goualher, G. L., Boomsma, D., Cannon, T., Kawashima, R., and Mazoyer, B. (2001). A probabilistic atlas and reference system for the human brain: International consortium for brain mapping (icbm). Philos Trans R Soc Lond B Biol Sci, 356(1412):1293–1322.

McLachlan, G. and Peel, D. (2000). Finite mixture models. Wiley Series in Probability and Statistics. Wiley-Interscience, first edition.

Mueller, S., Wang, D., Fox, M. D., Yeo, B., Sepulcre, J., Sabuncu, M. R., Shafee, R., Lu, J., and Liu, H. (2012). Individual variability in functional connectivity architecture of the human brain. Neuron, 77:586–595.

Munkres, J. (1957). Algorithms for the assignment and transportation problems. J Soc Ind Appl Math, 5(1):32–38.

Nummenmaa, L., Glerean, E., Viinikainen, M., Jääskeläinen, I. P., Hari, R., and Sams, M. (2012a). Emotions promote social interaction by synchronizing brain activity across individuals. Proc Natl Acad Sci, 109(24):9599–9604.

Nummenmaa, L., Glerean, E., Viinikainen, M., Jääskeläinen, I. P., Hari, R., and Sams, M. (2012b). Emotions promote social interaction by synchronizing brain activity across individuals. Proc Natl Acad Sci, 109(24):9599–9604.

Pajula, J., Kauppi, J.-P., and Tohka, J. (2012). Inter-subject correlation in fMRI: method validation against stimulus-model based analysis. PLOS ONE, 7(8):e41196.

Pajula, J. and Tohka, J. (2014). Effects of spatial smoothing on inter-subject correlation based analysis of fMRI. Magn Reson Imaging, 32(9):1114–1124.

Pamilo, S., Malinen, S., Hlushchuk, Y., Seppä, M., Tikka, P., and Hari, R. (2012). Functional subdivision of group-ICA results of fMRI data collected during cinema viewing. PLOS ONE, 7(7):e42000.

Reason, M., Jola, C., Kay, R., Reynolds, D., Kauppi, J.-P., Grobras, M.-H., Tohka, J., and Pollick, F. E. (2016). Spectators’ aesthetic experience of sound and movement in dance performance: A transdisciplinary investigation. Psychol Aesthet Creat Arts, 10(1):42.

Sabuncu, M. R., Singer, B. D., Conroy, B., Bryan, R. E., Ramadge, P. J., and Haxby, J. V. (2010). Function-based intersubject alignment of human cortical anatomy. Cereb Cortex, 20(1):130–140.

Salmi, J., Roine, U., Glerean, E., Lahnakoski, J., Nieminen-von Wendt, T., Tani, P., Leppämäki, S., Nummenmaa, L., Jääskeläinen, I., Carlson, S., et al. (2013). The brains of high functioning autistic individuals do not synchronize with those of others. NeuroImage: Clinical, 3:489–497.

Schwarz, G. et al. (1978). Estimating the dimension of a model. Ann Stat, 6(2):461–464.

Smith, D. V., Utevsky, A. V., Bland, A. R., Clement, N., Clithero, J. A., Harsch, A. E., Carter, R. M., and Huettel, S. A. (2014). Characterizing individual differences in functional connectivity using dual-regression and seed-based approaches. NeuroImage, 95:1–12.

Smith, J. (Accessed 02.05.2012). Spectral audio signal processing. http://ccrma.stanford.edu/˜jos/sasp/. online book.

Tan, P., Steinbach, M., and Kumar, V. (2014). Introduction to data mining. Addison-Wesley.

Tanskanen, T., Näsänen, R., Ojanpää, H., and Hari, R. (2007). Face recognition and cortical responses: effect of stimulus duration. NeuroImage, 35(4):1636–1644.

Thirion, B., Varoquaux, G., Dohmatob, E., and Poline, J.-B. (2014). Which fMRI clustering gives good brain parcellations? Front Neurosci, 8.

Tolhurst, D., Movshon, J. A., and Thompson, I. (1981). The dependence of response amplitude and variance of cat visual cortical neurones on stimulus contrast. Exp Brain Res, 41(3-4):414–419.

Trost, W., Frühholz, S., Cochrane, T., Cojan, Y., and Vuilleumier, P. (2015). Temporal dynamics of musical emotions examined through intersubject synchrony of brain activity. Soc Cogn Affect Neurosci, 10(12):1705–1721.

van den Heuvel, M., Mandl, R., and Hulshoff Pol, H. (2008). Normalized cut group clustering of resting-state fMRI data. PLOS ONE, 3(4):e2001.

Wang, D., Buckner, R. L., Fox, M. D., Holt, D. J., Holmes, A. J., Stoecklein, S., Langs, G., Pan, R., Qian, T., Li, K., et al. (2015). Parcellating cortical functional networks in individuals. Nature Neurosci, 18:1853–1860.

Wang, D. and Liu, H. (2014). Functional connectivity architecture of the human brain not all the same. Neuroscientist, 20(5):432–438.

Ward, J. (1963). Hierarchical grouping to optimize an objective function. J Am Stat Ass, 58(301):236–244.

Wilson, S. M., Molnar-Szakacs, I., and Iacoboni, M. (2008). Beyond superior temporal cortex: Intersubject correlations in narrative speech comprehension. Cereb Cortex, 18(1):230–242.

Xu, L. and Jordan, M. I. (1996). On convergence properties of the EM algorithm for Gaussian mixtures. Neural Comput, 8(1):129–151.

Zilles, K. and Amunts, K. (2013). Individual variability is not noise. Trends Cogn Sci, 17(4):153–155.

